# Refining Brain Stimulation Therapies: An Active Learning Approach to Personalization

**DOI:** 10.1101/2024.09.02.610880

**Authors:** Mohammad S. E. Sendi, Eric R. Cole, Brigitte Piallat, Charles A. Ellis, Thomas E. Eggers, Nealen G. Laxpati, Babak Mahmoudi, Claire-Anne Gutekunst, Annaelle Devergnas, Helen Mayberg, Robert E. Gross, Vince D. Calhoun

**Affiliations:** Wallace H. Coulter Department of Biomedical Engineering, Georgia Institute of Technology and Emory University, Atlanta, GA, United States; Department of Electrical and Computer Engineering at Georgia Institute of Technology, Atlanta, GA, United States; Tri-institutional Center for Translational Research in Neuroimaging and Data Science, Georgia State University, Atlanta, GA, United States; Current affiliation: Harvard Medical School and McLean Hospital, Boston, MA, United States; Department of Neurosurgery, Emory University School of Medicine, Atlanta, GA, 30322, United States; Univ. Grenoble Alpes, Inserm, U1216, Grenoble Institute of Neurosciences, Grenoble, France; Department of Biomedical Informatics, Emory University School of Medicine, Atlanta, GA, United States; Department of Neurology, Emory University School of Medicine, Atlanta, GA, 30322, United States; Emory National Primate Research Center, Atlanta, GA, 30322, United States; Departments of Neurology, Neurosurgery, Psychiatry and Neuroscience, Center for Advanced Circuit Therapeutics, Icahn School of Medicine at Mount Sinai, New York, NY, United States; Current affiliation: Department of Neurosurgery, Robert Wood Johnson Medical School, Rutgers University, New Brunswick, NJ, United States; Departments of Psychology and Computer Science, Georgia State University, Atlanta, GA, United States

## Abstract

Brain stimulation holds promise for treating brain disorders, but personalizing therapy remains challenging. Effective treatment requires establishing a functional link between stimulation parameters and brain response, yet traditional methods like random sampling (RS) are inefficient and costly. To overcome this, we developed an active learning (AL) framework that identifies optimal relationships between stimulation parameters and brain response with fewer experiments. We validated this framework through three experiments: (1) in silico modeling with synthetic data from a Parkinson’s disease model, (2) in silico modeling with real data from a non-human primate, and (3) in vivo modeling with a real-time rat optogenetic stimulation experiment. In each experiment, we compared AL models to RS models, using various query strategies and stimulation parameters (amplitude, frequency, pulse width). AL models consistently outperformed RS models, achieving lower error on unseen test data in silico (*p*<0.0056, *N*=1,000) and in vivo (*p*=0.0036, *N*=20). This approach represents a significant advancement in brain stimulation, potentially improving both research and clinical applications by making them more efficient and effective. Our findings suggest that AL can substantially reduce the cost and time required for developing personalized brain stimulation therapies, paving the way for more effective and accessible treatments for brain disorders.

## Introduction

Brain disorders comprise more than 600 conditions that impact an estimated million people worldwide every year^1^. These disorders cause impairment in the functionality of the central and peripheral nervous systems and lead to chronic physical, cognitive, and emotional disability. Deep brain stimulation (DBS), the focused delivery of electric current (usually a square pulse train) to the brain for the purpose of affecting neural or physiological processes, has seen increasing clinical use, and as of 2021, DBS had been used to treat more than 208,000 patients worldwide with a variety of neurological and neuropsychiatric disorders^2–6^. DBS has numerous benefits. (1) For example, relative to other techniques requiring surgical intervention, DBS is minimally invasive, does not damage brain tissue, and can be reversed. Moreover, (2) DBS parameters can be quickly and iteratively explored to maximize efficacy while minimizing side effects, and (3) DBS directly connects with the circuit pathophysiology that underlies overt symptoms ^7^.

Despite its clinical application in neurological and neuropsychiatric disorder treatment, the efficiency and consistency of DBS are still unsatisfactory, with highly variable results from patient to patient, which undermines its scalability and broader use ^8^. This challenge is twofold: (1) stimulation parameters must be evaluated in a systematic way using a data-driven approach and (2) disease- and patient-specific variables must be incorporated. The first step toward enabling more consistent, effective, and individualized DBS therapies is identifying diagnostic or therapeutic biomarkers that DBS can target or modulate. For example, in Parkinson’s disease (PD) in which the primary marker is known, algorithms can be developed to tune DBS parameters to achieve optimal outcomes, even though patient-specific factors such as ongoing degeneration may confound results. Moreover, the dynamics of different diseases vary significantly—epilepsy differs from PD, which - in turn - differs from depression. Therefore, a flexible platform to explore these dynamics with appropriate data is essential. Once those target response biomarkers are identified, although likely unique for each clinical indication, DBS systems can be used to make evidenced-based adjustments of stimulation parameters as needed for individual patients. However, finding the best set of stimulation parameters poses a key challenge. To find the best set of stimulation parameters for each individual, we need to understand how the brain, as a complex system, responds to various parameter sets. In other words, we need to learn a model between stimulation parameters and brain response (i.e., target response biomarkers) that is specific to a given disease. Modeling this link requires many samples, which is very costly and challenging and thus infeasible in the clinical setting. As such, an optimal experimental design for the identification of the samples that are most informative for uncovering a functional link is needed for both clinical and experimental settings ^9,10^.

Active learning (AL) is a machine learning method that can be used to address the optimal sampling problem. This process can help predictive models achieve higher performance with a limited number of samples by querying label (i.e., brain responses to particular sets of stimulation parameters) from an oracle (e.g., a human annotator or an experimental setup) for subset of informative instances ^11,12^. In the AL cycle, the learner starts with a small number of labeled samples. Next, the learner trains a model (e.g., for regression or classification) using the labeled samples, tests the model on the unlabeled samples, queries labels based on the informativeness of the unlabeled data, and then uses the updated knowledge to retrain a new model. AL evaluates the informativeness of unlabeled samples using a variety of query algorithms.

In this work, we present a novel AL framework developed to optimize (i.e., minimize the number of samples) an experiment modeling the relationship between stimulation parameters and brain response. We first developed a procedure for validating the utility of AL versus random sampling (RS) through *in silico* modeling with both synthetic and real data. Next, we developed a real-time AL procedure and validated the new approach in a real-time *in vivo* experimental setting. To demonstrate the generalizability of our method, we applied our AL framework to multiple datasets from different species. These included synthetic data generated from a Parkinson’s disease computational model of the basal ganglia circuit, real data collected from an epilepsy model in the non-human primate hippocampus, and optogenetic stimulation data collected from the rat hippocampus. By applying our approach across these diverse datasets and species, we aimed to highlight its versatility and potential for broad application in understanding and optimizing DBS therapies.

## Results

### Active learning outperforms random sampling in an *in silico* modeling experiment

We first developed a simulation procedure for the validation of the AL framework using synthetic data that consisted of three main phases (**Fig. 1a**). (1) We separated the data, containing stimulation parameters and their associated brain responses, into an unseen test dataset and training pool dataset – the set of all training data from which individual samples were iteratively drawn to build the model-specific training dataset. (2) We randomly selected a few samples from the training pool dataset to initiate a regression model between the stimulation parameters and their associated brain response (Phase 1 in **Fig. 1a**). (3) We added more samples, one at a time, to the model’s training dataset to iteratively improve the model. We used both RS and AL with different query strategies to determine which sample to add at each iteration (Phase2 in **Fig. 1a**). At each iteration of Phase 2, we evaluated the performance of the resulting model on the unseen test data. When evaluating model performance, we used the root mean square error (RMSE) between the actual and the predicted brain response (Phase3 in **Fig. 1a**). We repeated the entire process 1000 times for both AL and RS approaches, and we calculated the area under the curve (AUC) of RMSE, called AUC_RMSE after each iteration (**Fig. 1b**).

**Fig. 1.**
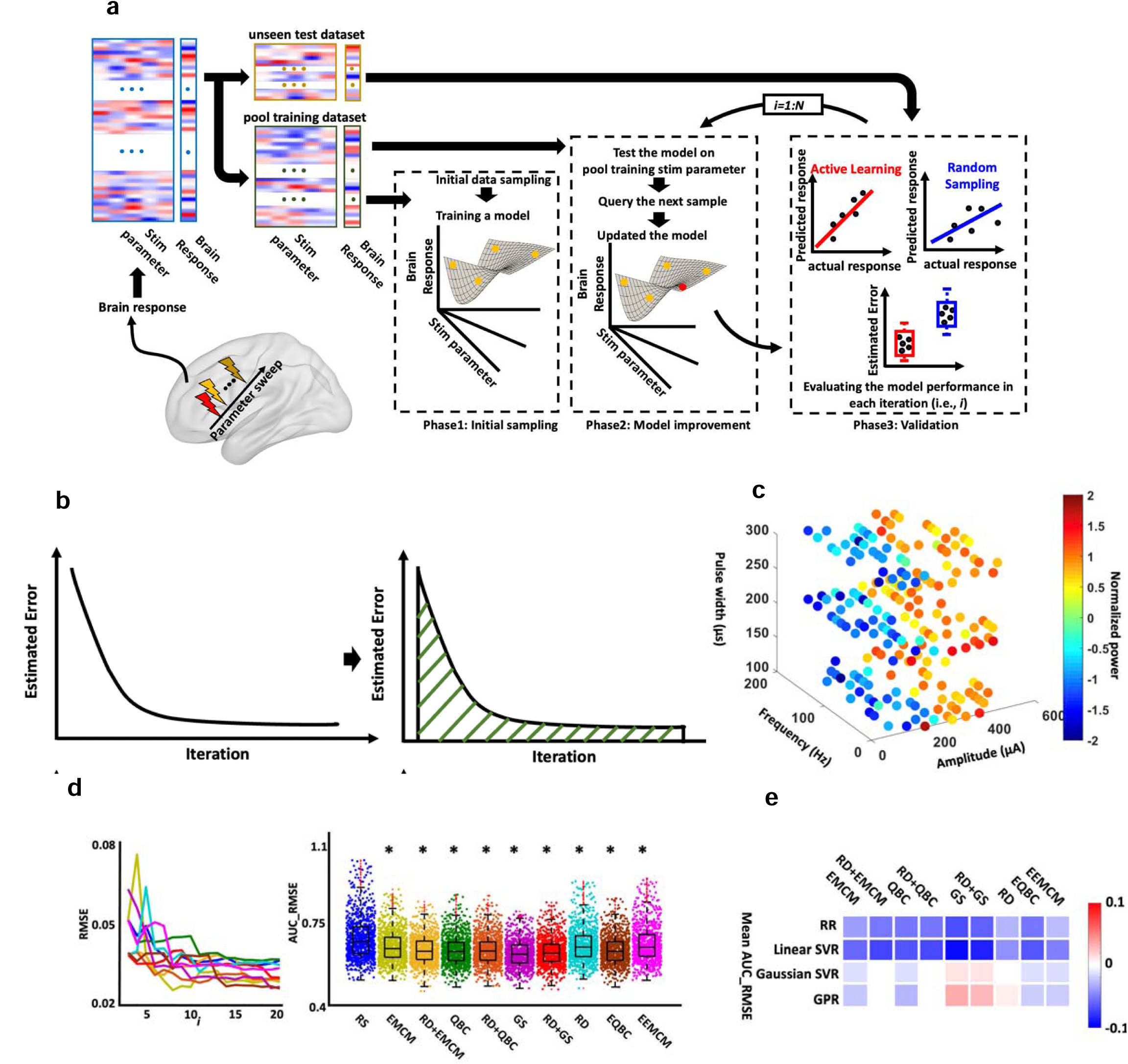
Active learning *in silico* modeling. **a)** Overview of the proposed stimulation protocol for validating the active learning (AL) framework. The procedure consists of three phases. In Phase 1, called *initial sampling*, a model is built between the response biomarker and stimulation parameters using a small subset of samples from the training dataset. In Phase 2, called *model improvement*, the model is iteratively updated by querying new samples, with a total of *N* samples collected. After each update, the model’s performance is evaluated on an unseen test dataset by estimating the Root Mean Square Error (RMSE) in Phase 3, referred as *validation*. **b)** At the end of the process, the area under the curve for RMSE, denoted as AUC_RMSE, is estimated. This entire process is repeated 1,000 times. Finally, the AUC_RMSE of the AL approach is compared with the AUC_RMSE obtained by Random Sampling (RS), and AL-based models that outperform the RS-based models by showing lower AUC_RMSE are identified after applying Bonferroni correction in two-sided two sample t-test of comparing AL and RS models performance (*p*<0.0056, *N*=1,000). **c)** The synthetic data generated for cortical-basal ganglia-thalamus network for PD model. **d)** Ridge regression (RR) from RS vs. AL with different query strategies on the cortex-basal ganglia-thalamus network PD model. Left panel: a representative trajectory of RMSE with the different number of samples queried through RS and AL with nine query strategies. Right panel: Mean RMSE_AUC of running 1,000 times with different AL query strategies and RS sampling. * represents those AL strategies having significantly lower mean AUC_RMSE than RS after Bonferroni correction in two-sided two sample t-test of comparing AL and RS models performance (*p*<0.0056, *N*=1,000). **e)** A summary plot comparing AL-based model versus RS-based model performance on unseen test dataset for different regression models and query strategies in two-sided two sample t-test of comparing AL and RS models performance (*p*<0.0056, *N*=1,000). This plot illustrates the difference in AUC_RMSE between models obtained using AL and those obtained using RS for each regression model. In this figure, colored cells highlight results that remained significant after applying Bonferroni correction.

To validate the proposed AL framework through the simulation process, we generated synthetic data using a biophysical model of the cortex-basal ganglia-thalamus network in a 6-OHDA lesioned rat with Parkinson’s disease (See **Supplementary Fig.1**) ^13^. We stimulated the subthalamic nucleus (STN) and swept amplitude, frequency, and pulse width while estimating the globus pallidus internus (GPi) beta (13-30 Hz) power for each stimulation parameter. This resulted in 200 different samples (see first dataset in Methods, **Fig. 1c**, and **Supplementary Fig.2**). We used 20% (i.e., 40 samples) as unseen test data and 80% (i.e., 160 samples) as a training pool dataset. We next trained a ridge regression (RR) model between STN stimulation parameters and GPi beta power with both AL and RS approaches in which we started with three initial samples and added more samples from the training pool dataset until we had 20 training samples. As we expected, the model performance, regardless of sampling strategy, improved with the addition of more samples to the model training data (**Fig. 1d** left panel). Our results showed that all query strategies in AL had significantly lower mean AUC_RMSE on unseen datasets compared to the RS method, as determined by a two-sample t-test comparing the AUC_RMSE of each AL method against that of the RS model. (**Fig. 1d** right panel, * represents Bonferroni corrected *p*<0.0056).

We also trained support vector regression (SVR) models with linear and Gaussian kernel functions and Gaussian process regression (GPR) models using both RS and AL approaches and found that better performance resulted from the AL approach relative to RS (**Fig. 1e**, see **Supplementary Fig.3, 4, & 5**). **Fig. 1e** shows the difference between the AUC_RMSE of the model performance obtained by AL and the AUC_RMSE obtained by RS for each regression model. The results shown in **Fig. 1e** suggest that the AL approach can provide a better model than the RS approach. However, the query strategy and regression model must be selected carefully. For example, representativeness and diversity (RD) combined with query by committee (QBC) provided a better result than RS in RR and linear SVR, while it did not show a model performance improvement in Gaussian SVR and GPR based on their AUC_RMSE metric (**Fig. 1e**).

### Active learning results reproduced across experiments within a subject

The previous section proposed the AL framework for designing an optimal experiment to get the best model between the simulation parameters and brain response. After validating the advantages of the AL vs. RS approach, we next questioned whether the results were replicable across multiple sessions of the parameter sweep data within a subject. Answering this question was necessary for validating the robustness of the proposed AL framework. To answer this question, we designed a parameter sweep experiment, stimulating the hippocampal network of a non-human primate (NHP) while recording from the same network using the Summit RC+S (Medtronic, Minneapolis, USA) as shown in **Fig. 2a** (see second dataset in *Methods*) and quantifying the post-stimulation theta power. Asynchronous distributed multielectrode stimulation or ADMES, which has been shown to reduce seizure frequency in rodent models ^14^, was used to stimulate the NHP hippocampus (**Fig. 2b**)^15^. We repeated the same experiment four times and recorded 100 samples. Next, we ran the AL vs. RS *in silico* modeling in each session to validate the reproducibility of the results. We had 20 unseen samples and 80 training pool samples in this simulation. Finally, we compared the mean AUC_RMSE of RS with those of AL with different query strategies while training a regression model between the stimulation parameters and post-stimulation theta power.

**Fig. 2.**
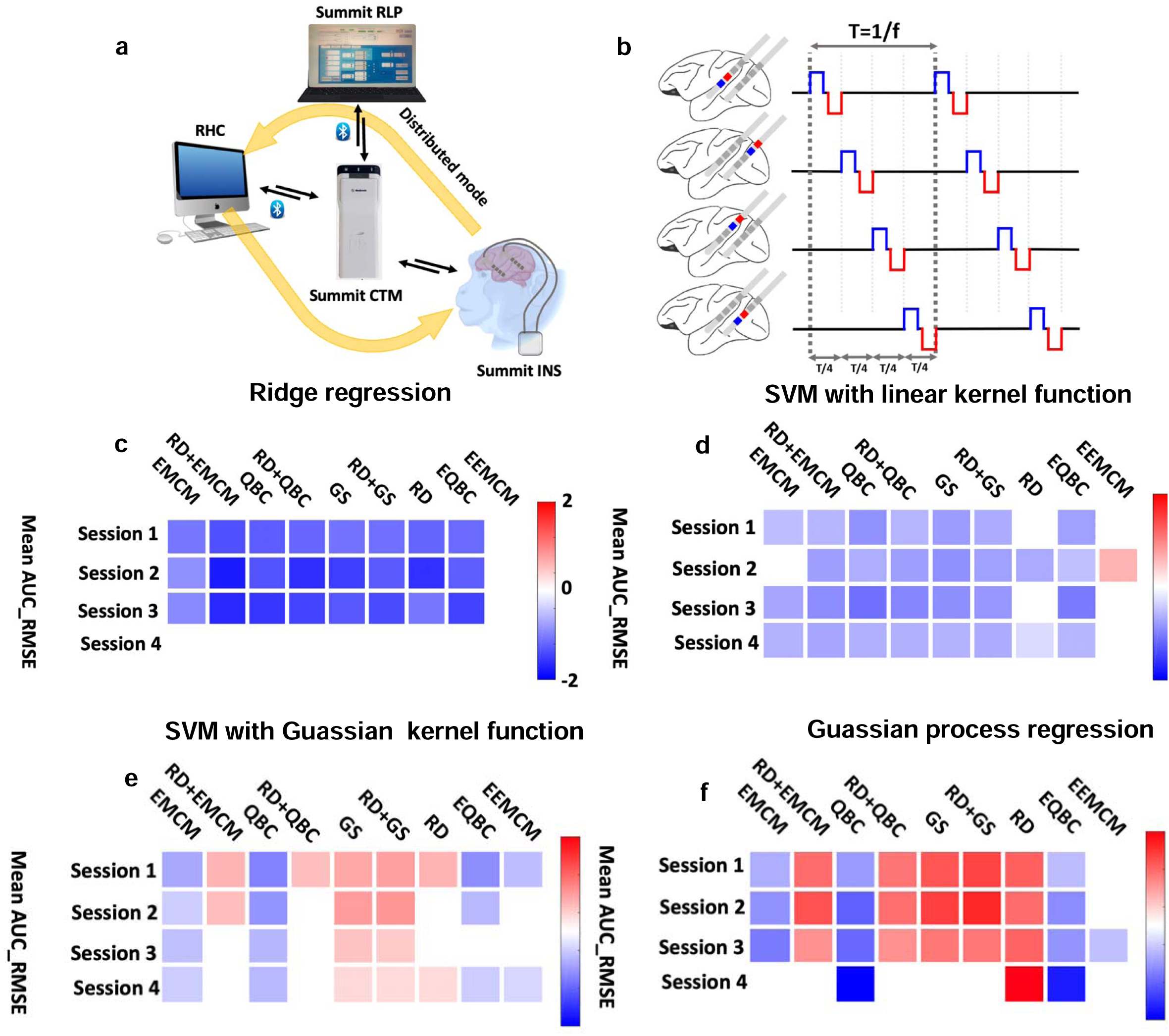
Reproducibility of active learning within subject. **a)** Summit implantable neurostimulator (INS) can have bidirectional communication with the researcher’s host computer (RHC) and research lab programmer (RLP) via the clinician telemetry module (CTM). With the RLP, we can set the safety level of stimulation parameters, and with the RHC, we can control the INS configuration for real-time recording and stimulation. Two leads contain eight stimulation and recording sites implanted in the hippocampus of a non-human primate. **b)** Conceptual schematic of the asynchronous distributed multielectrode stimulation (ADMES). **c)** A summary plot comparing performance of AL-based models versus RS-based models on an unseen test dataset for a ridge regression (RR) model with different query strategies in which we repeated the simulation process for each model 1,000 times. This panel illustrates the difference in AUC_RMSE between models obtained using AL with various query strategies and those obtained using RS. In this figure, colored cells highlight results that remained significant after applying the Bonferroni correction in two-sided two sample t-test of comparing AL and RS models performance (*p*<0.0056, *N*=1,000). The model obtained from all query strategies except EEMCM showed better AUC_RMSE compared with the RS-based model in all sessions except session 4. **d)** A summary plot comparing the performance of AL-based models versus RS-based models on an unseen test dataset for a support vector regression model with linear kernel function model with different query strategies. The plot illustrates the difference in AUC_RMSE between models obtained using AL with various query strategies and those obtained using RS. In this figure, colored cells highlight results that remained significant after applying the Bonferroni correction in two-sided two sample t-test of comparing AL and RS models performance (*p*<0.0056, *N*=1,000). We observed an improvement in the regression model performance using AL in all sessions when we used AUC_RMSE as a metric. **e)** A summary plot comparing the performance of AL-based models versus RS-based models on an unseen test dataset for a support vector regression model with Gaussian kernel function model with different query strategies. This plot illustrates the difference in AUC_RMSE between models obtained using AL with various query strategies and those obtained using RS. In this figure, colored cells highlight results that remained significant after applying the Bonferroni correction in two-sided two sample t-test of comparing AL and RS models performance (*p*<0.0056, *N*=1,000). We observed an improvement in the regression model AUC_RMSE using EMCM and QBC in all sessions. **f)** A summary plot comparing the performance of AL-based models versus RS-based models on an unseen test dataset for Gaussian process regression (GPR) with different query strategies. This plot illustrates the difference in AUC_RMSE between models obtained using AL with various query strategies and those obtained using RS. In this figure, colored cells highlight results that remained significant after applying the Bonferroni correction in two-sided two sample t-test of comparing AL and RS models performance (*p*<0.0056, *N*=1,000). We observed an improvement in the regression model AUC_RMSE using QBC and EQBC in all sessions.

A summary plot presented in **Fig. 2c** illustrates the performance comparison of AL-based RR models with RS-based RR models across all 1000 iterations on unseen test data for four sessions. In this plot, each cell represents the difference in mean AUC_RMSE between the AL-based model and the corresponding RS-based model. Colored boxes that are highlighted indicate significant results between the AL and RS model performance, as determined by Bonferroni comparisons (*p* < 0.0056). As shown in **Fig. 2c**, all AL query strategies enhanced model performance more than the RS approach in sessions 1, 2, and 3. However, none of the query strategies surpassed the RS approach in session 4 when assessing model performance with mean AUC_RMSE.

**Fig. 2d** presents a similar summary plot for SVR with a linear kernel function. Based on the AUC_RMSE metric, the models obtained with seven of the nine query strategies outperformed the corresponding RS model on the unseen test data of each session. **Fig. 2e** presents the summary plot for SVR with a Gaussian kernel function. According to the AUC_RMSE metric, the model obtained by expected model change maximization (EMCM) and QBC outperformed the corresponding RS-based model in all four sessions. Additionally, the model from enhanced QBC (EQBC) outperformed the RS-based model in three sessions, and the model from enhanced EMCM (EEMCM) outperformed the RS-based model in two sessions.

**Fig. 2f** presents the summary plot for GPR models. According to the AUC_RMSE metric, the model obtained by QBC and EQBC outperformed the corresponding RS-based model in all four sessions. Overall, our observations confirmed the superiority of the AL approach over the RS approach, as demonstrated by its replicability across multiple sessions within a single subject dataset. This consistency across sessions underscores the robustness of our AL framework and its potential generalizability to different experimental scenarios, as demonstrated here within the NHP model.

### Dynamic modeling makes the active learning sampling efficiency more pronounced relative to random sampling

Another question was whether brain dynamics would alter the performance of our model. Unlike many other AL regression applications, in which the model between input and output is static and does not change across experiments, the brain is highly dynamic. The brain response to stimulation could differ depending on the pre-stimulation brain state (**Fig. 3a**). For example, a recent study showed that direct stimulation enhanced memory only when stimulation was applied during brain states linked with poor memory outcomes ^15^. Therefore, considering the pre-stimulation brain state while modeling functional links between stimulation parameters and post-stimulation brain response may be necessary to have a more accurate model. To address this question, we expanded the framework introduced in **Fig. 1a** to consider brain dynamics by adding the pre-stimulation brain state (called *brain state dynamic modeling*). We used one session of the NHP dataset in which we calculated the pre-stimulation theta power and added that to the model. Similar to the previous section, we used 80 samples (80% of the whole dataset) for the training pool dataset and 20 samples (20% of the whole dataset) for the unseen test data.

**Fig. 3.**
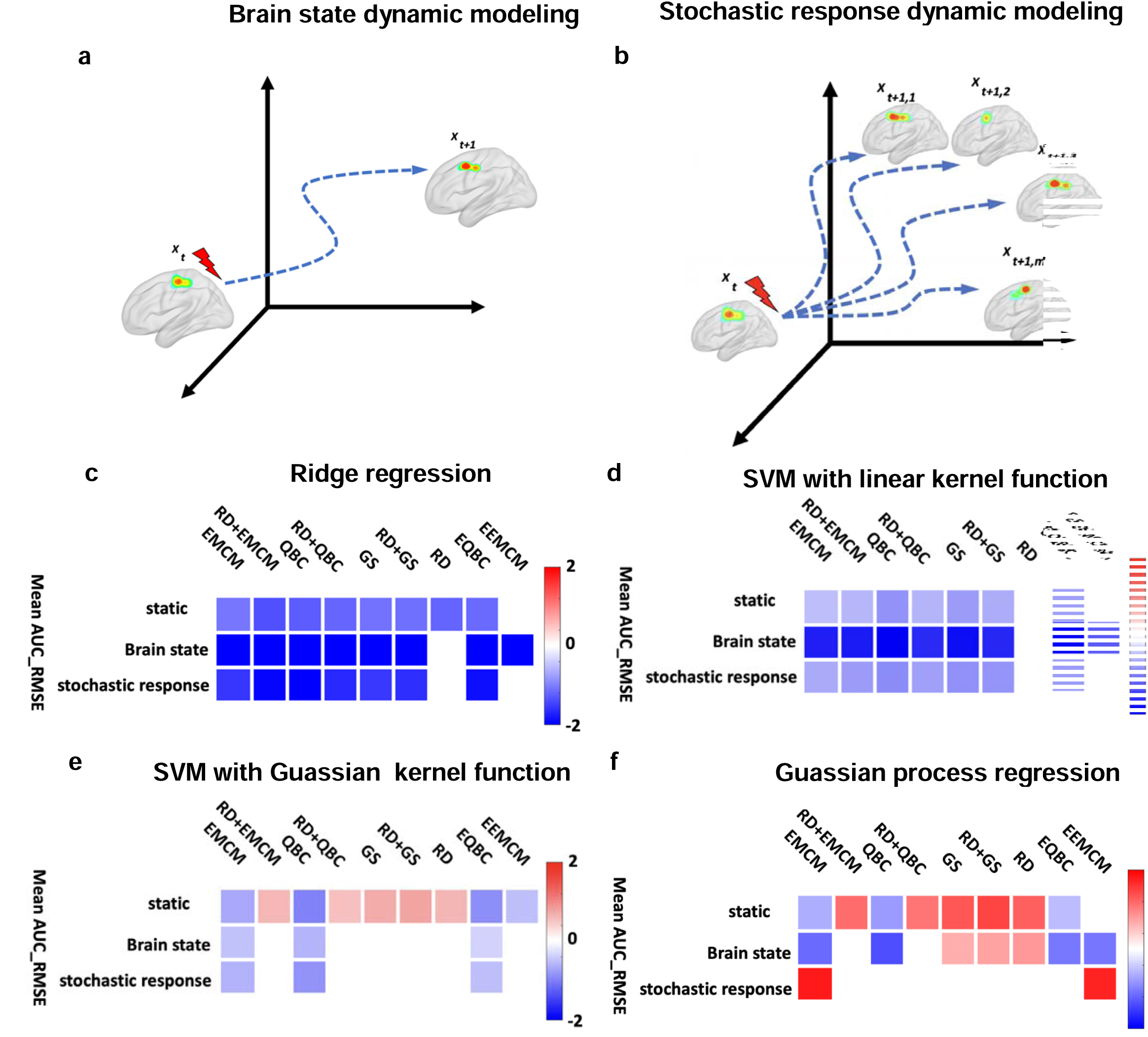
Dynamic modelling. **a)** *Brain state dynamic modeling*: example trajectory through state space in a dynamic model. Incorporating the state of the brain prior to the stimulation in the dynamic model is the only difference between the dynamic and static models. **b)** *Stochastic response dynamic modeling*: In contrast to other systems, the brain is dynamic in that the system response associated with an input set is different across multiple system identification tests. That means the brain response is different when we administrate the same stimulation parameters. **c)** A summary plot comparing the performance of AL-based models versus RS-based models on an unseen test dataset for a ridge regression model with different query strategies in *static*, *brain state dynamic modeling*, and *stochastic response dynamic modeling*. This graph illustrates the difference in AUC_RMSE between models obtained using AL with various query strategies and those obtained using RS in which we repeated the simulation process for each model 1000 times. In this figure, colored cells highlight results that remained significant after applying the Bonferroni correction (*p*<0.0056, N=1,000). In the *static* setting for the RR model, eight of nine query strategies outperformed the RS approach. A similar trend was observed in *brain state dynamic modeling*, with eight out of nine strategies surpassing RS performance in the unseen test data. In *stochastic response dynamic modeling*, seven out of nine strategies outperformed RS. The AL models consistently outperformed the RS models, with a greater suppression of AUC_RMSE in *brain state dynamic modeling* and *stochastic response dynamic modeling* settings. **d)** A summary plot comparing the performance of AL-based models versus RS-based models on an unseen test dataset for a support vector regression model with linear kernel function model with different query strategies in *static*, *brain state dynamic modeling*, and *stochastic response dynamic modeling* in which we repeated the simulation process for each model 1000 times. Seven out of nine AL approaches in both the *static* and *stochastic response dynamic modeling* settings, and eight out of nine in the *brain state dynamic modeling* setting, outperformed the RS model on the unseen test dataset. The improvement is more pronounced in the *brain state dynamic modeling* setting. **e)** A summary plot comparing the performance of AL-based models versus RS-based models on an unseen test dataset for a support vector regression model with Gaussian kernel function model with different query strategies in *static*, *brain state dynamic modeling*, and *stochastic response dynamic modeling*. In the static scenario, five AL-based models exhibited higher AUC-RMSE than the RS model. However, in both the *brain state dynamic modeling* and *stochastic response dynamic modeling* scenarios, none of the AL-based models showed higher AUC-RMSE than the RS model. **f)** A summary plot comparing the performance of AL-based models versus RS-based models on an unseen test dataset for Gaussian process regression (GPR) with different query strategies in *static*, *brain state dynamic modeling*, and *stochastic response dynamic modeling* in which we repeated the simulation process for each model 1000 times. In the static scenario, five AL-based models exhibited higher AUC-RMSE compared to the RS model. However, in *brain state dynamic modeling* none of the AL-based models showed higher AUC-RMSE than the RS model.

Additionally, in contrast to other systems, the brain system response associated with a specific input is stochastic in that its response is not identical across repeated samples. In practice, this implies that identical stimulation parameters administered at different points in time can yield different brain responses (see **Fig. 3b**). To model this aspect of brain dynamics, we developed a new approach and updated our original framework to use multiple responses of the same parameter set collected across multiple sessions of the parameter sweep experiment (called *stochastic response dynamic modeling*). Similar to the original framework (i.e., **Fig. 1a**), the entire simulation process of the new framework contained three phases (see **Supplementary Fig.10**). First, we separate the entire dataset (i.e., stimulation parameter sets and their associated brain response) into the unseen validation and training pool dataset. Then in Phase 1, we randomly selected three stimulation parameters and their associated brain responses (one from *m* available responses for each stimulation parameter). In Phase 2, we iteratively improved the model by increasing the training data size across *N* iterations. In more detail, based on the AL or RS approach recommendation, we selected one stimulation parameter and its associated brain response (one from all *m* available samples) and updated the model. After updating the model, we validated the model on the unseen validation dataset and calculated the RMSE between the predicted and actual brain response (Phase 3). We calculated the area under the curve or AUC of the RMSE at the end of the process. We ran the entire process 1000 times to ensure statistical significance of results. Finally, we compared the RS-model’s mean AUC_RMSE with the AL-model’s performance with all query strategies. We used the NHP dataset (second dataset in *Methods*), which includes four different post-stimulation brain responses for each of 100 different stimulation parameter sets (i.e., *m*=4). Again, we used 80 samples of the entire dataset as the training pool and 20 samples of the unseen test data while training a regression model between the stimulation parameters and hippocampal theta power.

**Fig. 3c** presents a summary plot of the models’ performance, specifically showing the difference in mean AUC_RMSE between the active learning approach and the corresponding RS-based model for RR models with various query strategies in static, *brain state dynamic modeling*, and *stochastic response dynamic modeling* settings. For the RR model in the *static* setting, it was observed that eight out of nine query strategies surpassed the performance of the RS approach. In *brain state dynamic modeling*, a similar trend was observed, with eight of nine query strategies outperforming the model obtained through RS in the unseen test data. Additionally, in the *stochastic response dynamic modeling* setting, the models obtained from seven of nine query strategies outperformed the model obtained through RS. Notably, the models obtained through AL consistently outperformed those obtained through RS, significantly reducing AUC_RMSE, particularly in the *brain state dynamic modeling* and *stochastic response dynamic modeling* settings.

**Fig. 3d** presents similar results for SVR with a linear kernel function. As depicted in the figure, seven of nine AL approaches in both the *static* and *stochastic response dynamic modeling* settings and eight of nine AL approaches in the *brain state dynamic modeling* setting outperformed the model obtained through RS on the unseen test dataset. However, the extent of AUC_RMSE reduction of AL relative to RS is more pronounced in the *brain state dynamic modeling* setting.

In **Fig. 3e**, which presents the results for SVR with a Gaussian kernel function, five AL-based models exhibited higher AUC-RMSE compared to the model obtained through RS in the *static* scenario. However, none of the AL-based models showed higher AUC-RMSE than the RS model in the *brain state dynamic modeling* and *stochastic response dynamic modeling* scenarios.

Additionally, as illustrated in **Fig. 3f** for GPR, three AL-based models outperformed the RS-based model on unseen test data in the *static* setting, four AL-based models outperformed the RS-based model in *brain state dynamic modeling*, and none of AL-based models outperformed the RS-based model in *stochastic response dynamic modeling*. Notably, the AUC-RMSE in AL-based models was lower compared to that in RS-based models in *brain state dynamic modeling* and *stochastic response dynamic modeling* scenarios when compared with the *static* scenario. In summary, our results demonstrate that the AL approach outperforms the RS approach in both static and dynamic models. However, we also show evidence that the AL approach is more beneficial when incorporating brain dynamics, as opposed to only considering a static brain response.

### *In silico* simulation for designing an *in vivo* real-time AL experiment

We next asked whether the AL-based model would outperform the RS-based model in a real-time *in vivo* experiment. To answer this question, we used data collected from a parameter sweep experiment in a rat optogenetic stimulation paradigm for *in silico* modeling and to select prior information before the *in vivo* real-time experiment (third dataset in *Methods*). We selected this paradigm due its well-characterized response and prior use as a model system for computational brain stimulation methods ^16,17^. A rectangular pulse train of light was delivered to the medial septum of an adult male Sprague-Dawley rat through the implanted fiber optic at all combinations of amplitude or intensity (between 10 to 50 mW mm^−2^), pulse-width (between 2 to 10 ms), and pulse frequency (between 5 to 42 Hz) for a total of 72 combinations of stimulation parameters. Stimulation parameters were applied in a random order for 5 s. The hippocampal LFP was recorded during stimulation, and slow gamma power was computed during stimulation to provide the target brain response for learning. All samples collected through this process are shown in **Fig. 4a**.

**Fig. 4.**
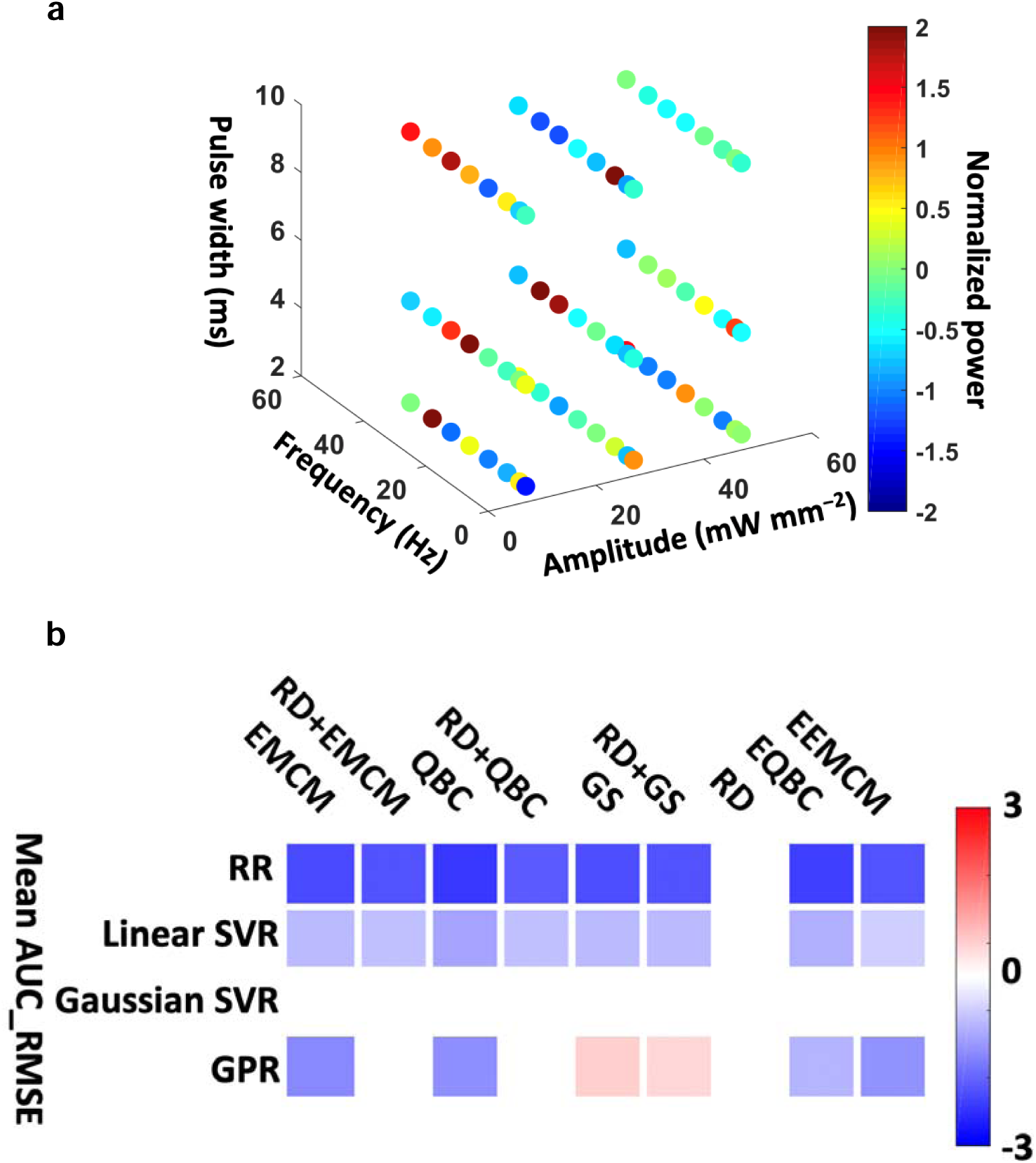
In silico modeling result before *in vivo* experiment. **a)** Normalized gamma power for different stimulation parameters. **b)** Summary plot of models’ performance on the medial septum optogenetic parameter sweep data. This graph illustrates the difference in AUC_RMSE between models obtained using AL with various query strategies and those obtained using RS in which we repeated the simulation process for each model 1000 times. In this figure, colored cells highlight results that remained significant after applying the Bonferroni correction in two-sided two sample t-test of comparing AL and RS models performance (*p*<0.0056, N=1,000). The ridge regression model with query by committee (QBC) achieved the lowest mean AUC_RMSE among all regression models and query strategies. Consequently, we selected this model for the *in vivo* experiment.

We used the framework proposed in **Fig. 1a** to compare the RS-model with the AL-models (with different query strategies) and to find the best query strategy and regression model for later *in vivo* deployment. To ensure that we had adequate samples to validate the model, we allocated 20 samples for validation in each iteration, while the remaining 52 samples were used to create the training pool dataset. We started with three initial samples (equal to the number of parameters), iteratively queried more samples (one at a time) and added those samples to the training dataset through both AL (with different query strategies) and RS. We estimated RMSE on the unseen datasets at each iteration. We repeated the simulation 1000 times and estimated the mean AUC for RMSE. By adding more samples in Phase2 (**Fig. 1a**), we obtained a better model with less RMSE on the unseen test data (**Fig. 4b**). The summary plot of all regression models and query approaches used on this dataset reveals the following insights: 1) AL-based models, except the one obtained through RD, outperformed RS-based models in RR and SVR with a linear kernel, as indicated by lower mean AUC_RMSE. 2) Among all query strategies, EMCM, QBC, EQBC, and EEMCM demonstrated lower mean AUC_RMSE in GPR models. 3) RR with QBC achieved the lowest mean AUC_RMSE among all regression models and query strategies. Based on these findings, we selected the RR model with QBC for the *in vivo* experiment.

### Active learning improves sample-efficiency relative to random sampling *in vivo*

After identifying the optimal regression model and query strategy, we conducted a real-time *in vivo* experiment to model the relationship between stimulation parameters and slow-gamma power using both RS and AL approaches (see **Fig. 5a** and *Methods*). We observed that the model derived from the QBC approach (represented by green circles) exhibited significantly lower mean AUC_RMSE compared to the model based on the RS approach (represented by blue circles) across 25 unseen test data points in each of 20 experimental sessions (*p*=0.0036, *N*=20), as illustrated in **Fig. 5b**. This outcome aligns with our *in silico* modeling results, where an improvement in AUC_RMSE was noted when the regression model was obtained through AL.

**Fig. 5.**
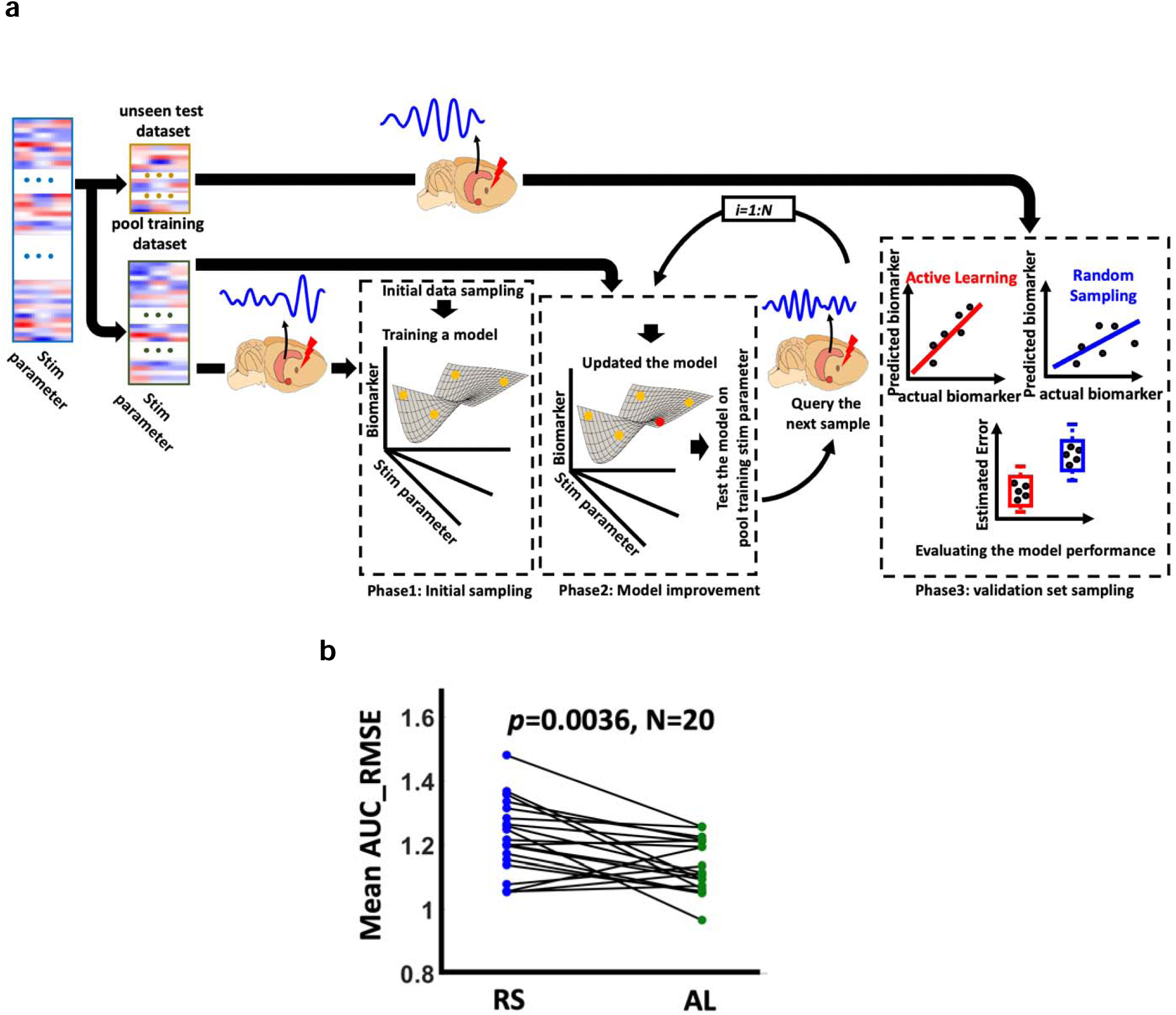
An overview of proposed *in vivo* active learning framework. a) The experiment contains three phases. *Phase1* referred as *initial sampling:* We assigned the entire parameters set (1080 samples) to the unseen test (25 samples) and training pool (1055 samples) sets. The 25 unseen test samples were not used in training phase. Next, we selected 3 samples from the training pool and simulated medial septum, recorded local field potential from CA1 during the stimulation, and estimated slow gamma power. Then, we trained an initial regression model (between the stimulation parameters and the slow-gamma power) based on three collected samples. *Phase2* referred as *model improvement:* We iteratively increased training samples size (one at a time). We added 72 samples through both active learning and random sampling process. *Phase3* referred as *validation set sampling:* We stimulated the medial septum with the stimulation parameters of the test dataset and recorded from CA1. We collected 25 samples in this phase. b) AL vs RS in a real-time *in vivo* medial septum optogenetic experiment by comparing the mean AUC_RMSE of the models on the unseen dataset. We found the ridge regression (RR) model with query by committee or QBC (green circles) outperformed the model from random sampling or RS (blue circles) on 25 unseen test datasets across 20 experimental sessions (*p*=0.0036, *N*=20).

## Discussion

Modern control techniques are based on a good understanding of the system (i.e., a model between input and output) to be controlled ^18^. This model can be based on physical laws. However, modeling a system as complex as the brain and its dynamics is very difficult. System identification refers to the mathematical modeling process of extracting information about a system from measured input-output data ^19^. For brain system identification, it is necessary to perturb input signals (i.e., the stimulation parameters) and observe the system output (i.e., the brain response). However, existing brain system identification approaches used in clinical settings apply unoptimized random sampling that is inefficient and costly.

Active learning, which is referred to as *experimental design* in statistics, is a subfield of machine learning and statistics that provides a smart solution for designing an experiment in which human decision-making is suboptimal for the task ^10^. The main rationale underpinning active learning is that data collection is costly, so query points should be selected such that the performance of the model being identified is optimized. More specifically, active learning is a paradigm in which machine learning models direct the learning process by providing dynamic suggestions/queries for the *next-best experiment*.

Here, we developed an *in silico* simulation procedure and ran a stimulation parameter sweep using the model of the cortex-basal ganglia-thalamus network in an 6-OHDA lesioned rat with Parkinson’s disease ^20^. We generated synthetic data in which we swept STN stimulation parameters (i.e., stimulation amplitude, frequency, and pulse width) while estimating GPi beta power as a key biomarker of Parkinson’s diseases. We used this dataset to validate our AL framework by comparing the model obtained through RS and AL algorithms with nine different query strategies (see **Fig. 1e**). In this simulation, we showed that the models trained through the AL process outperform the RS-based model for both linear and nonlinear regression models. However, not all query strategies-built models that outperformed the RS-based model. This indicates that careful *in silico* modeling is needed to select the best regression model parameters and query strategy prior to conducting an *in vivo* experiment. This finding is similar to that of a previous study elucidating the regulatory relationships among genes and proteins in yeast that asserted that prior knowledge is needed before starting an experiment ^21^.

Previous studies have used Bayesian optimization (BaO) to optimize DBS experimental procedures. However, there are fundamental differences between AL and BaO in DBS. First, the objective of BaO is to select the best stimulation parameter and collect the most informative observation to maximize or minimize the brain biomarker response, while minimizing the number of trials. For example, a recent study adapted BaO to suppress subthalamic nucleus (STN) beta power in a computational model of PD ^22^. In contrast, the AL objective is to find the optimal regression model between DBS parameters and brain response with a minimum number of trials. Once we have the most accurate model, we can predict the brain response to any untested DBS parameters throughout the defined input space (rather than only one optimal input). As such, this model can potentially be used to minimize or maximize the brain response by finding the best DBS parameters. Second, BaO is limited to use with Gaussian process regression (GPR), while AL is compatible with any kind of regression model. As we have shown, GPR might not be the best model for predicting the brain response from the DBS parameters. Therefore, even if AL had no other advantages over BaO, it would still have the potential to outperform the BaO approach simply by enabling the use of better regression models. A future study is needed to compare BaO and AL in minimizing or maximizing a brain response biomarker in the context of DBS.

After developing our AL approach, we next tried to answer two important questions prior to *in vivo* experimentation. The first question was whether AL outperforms RS across multiple sessions within a subject. To answer this question, we designed a parameter sweep experiment in which we collected four sessions of NHP data using the Summit RC+S device. Next, we compared AL to RS, with a different regression model, in each session separately. We found that AL outperformed the RS model in all sessions. However, within query strategies, results were not always consistent. For example, in the RR model, the RD approach outperformed the RS approach in sessions 1,2 and 3 but not session 4 (**Fig. 2c**).

We also explored how considering the brain dynamics changes AL and RS test results. We updated the proposed AL framework and added the pre-stimulation brain state, called *brain state dynamic modeling* (**Fig. 3a**). We also proposed a new approach to consider brain dynamics based on multiple data sessions, which we called *stochastic response dynamic modeling* (**Fig. 3b**). In *stochastic response dynamic modeling*, we randomly selected the brain response out of four sessions of the NHP dataset for a given stimulation parameter set. The main finding of comparing *static* with *brain state dynamic modeling* and *stochastic response dynamic modeling* is that adding dynamics to the model further demonstrates the utility of the AL approach over the RS one. For instance, in the RR model using the *static*, *brain state dynamic modeling*, and *stochastic response dynamic modeling* approaches, at least seven AL query strategies outperformed the RS approach. Notably, in the *brain state dynamic modeling* and *stochastic response dynamic modeling* approaches, the improvement in model performance by AL was significantly greater, as evidenced by a more substantial reduction in AUC_RMSE. This trend was also observed in other regression models.

We also demonstrated how *in silico* modeling informed the design of an *in vivo* experiment. We first collected a parameter sweep dataset using optogenetic stimulation of the medial septum in an anesthetized rat. Then, using the simulation process proposed in **Fig. 1a**, we compared the performance of the regression models, including RR, SVR with linear and Gaussian kernel functions, and GPR obtained from RS with those from AL one with different query strategies. This simulation process found that the RR model obtained from the QBC query strategies showed the best performance relative to other regression models and query strategies. Choosing the best regression model type that broadly captures the appropriate structure of the experimental data is vital for AL, as using the wrong regression model might mislead the real-time experiment ^11^.

Next, we used this prior knowledge and ran an *in vivo* experiment with the same setup we used for the *in silico* modeling. We ran 20 sessions of the experiment using the procedure shown in **Fig. 5a**. To the best of our knowledge, this is the first real-time implementation of the AL framework in a brain stimulation experiment, demonstrating its utility over RS in neuromodulation. After completing 20 sessions of the proposed experimental procedure, we compared the RS and AL with QBC approach through a post hoc analysis. We found that the model obtained by the AL approach outperformed the RS model based on RMSE metrics (**Fig. 5b**).

Although our results showed that the AL approach outperformed the RS one overall by having less mean AUC_RMSE, we did find that the RS approach outperformed AL in some sessions. The AL approach might have misled the experiment by querying the wrong data. Collecting noisy data might be a possible explanation for how AL misled the experiment. While we tried to minimize motion artifacts and behavioral state-dependent confounds with anesthesia, there may have been some noisy data at one iteration of the experiment that mislead the remainder of the AL data collection for some sessions. It is worth mentioning that querying noisy data would not be a problem in a conventional AL framework when working with a human oracle because we assume that human oracles never provide wrong information in the AL process ^23^. Additionally, to reduce computational intensity, we did not use cross-validation to optimize the regression model between the optogenetic parameters and slow-gamma power during the experiment. As such, the model used to query the next stimulation parameters is unoptimized, which could mislead the experiment.

There are a few limitations associated with our proposed framework. First, we only focused on traditional machine learning-based regression models. Future studies are needed to explore the performance of deep learning models obtained from random sampling and active learning^24^. Second, we only focused on single-output regression models, and future studies might explore multiple-output regression models^25^. Also, we validated the second dynamical modeling approach using only four sessions of the parameter sweep experiment, and further study is needed to explore the proposed framework with a greater number of sessions. We assume that having more sessions would further demonstrate the benefits of the AL approach over the RS one. In this study, we only focused on the reproducibility of AL versus RS test results for within-subject data, and future study is needed to explore the reproducibility of results across subjects. The data provided through the restoring active memory (RAM) project ^26^, in which the same parameter sweep experiment was done across multiple patients, could be a good resource to evaluate whether AL-based models would outperform RS-based models across patient data.

There are also a few limitations associated with our *in vivo* studies. First, we chose the regression model and query strategies based on data from only one parameter sweep. While we showed that the result of query strategies might be replicated across multiple sessions of the same parameters sweep experiment, a future study is needed to find the best regression model and query strategies based on data from multiple experiments. Noisy data collection could mislead the AL experiment, and if we consider noisy data to be outliers, other query strategies based on *representativeness* and *diversity* (RD) could potentially provide a solution. There is a line of work in the AL community that explicitly focuses on dealing with noisy labels ^27–29^. Future studies are needed to compare other AL approaches that could alleviate the effect of noisy labels in our experimental setup. Also, we ran the real-time experiment on only one animal. A future study is needed to evaluate the reproducibility of the real-time experiment across subjects.

In summary, we proposed an AL framework and developed a simulation procedure to validate its utility over the RS approach. We validated the reproducibility of the method across multiple sessions with the same subject and demonstrated the increased efficacy of the AL approach when accounting for brain dynamics in our dynamical models. Additionally, we showed how the *in silico* modeling can provide the prior information needed for a real-time *in vivo* experiment. Finally, we ran a real-time *in vivo* experiment of the AL and RS approach and showed that AL is more sample-efficient than RS.

## Methods

### Datasets

The current study comprises three datasets. The first dataset, referred to as the Synthetic Parkinson’s Disease (PD) model, involves synthetic data generated to validate the active learning (AL) framework. This validation utilized a biophysical model of the cortex-basal ganglia-thalamus network in a 6-OHDA lesioned rat, simulating Parkinson’s disease, as proposed in the referenced work (https://github.com/ModelDBRepository/206232)^13^. The network includes the cortex, striatum, and subthalamic Huxley-type neurons (refer to Supplementary Material for details). To simulate both healthy control (HC) and PD models, we adjusted the M-type potassium current for both the direct and indirect medium spiny neurons (MSNs) by setting the maximal conductance at 2.6 mS/cm² and 1.5 mS/cm², respectively. We varied the subthalamic nucleus DBS amplitude, frequency, and pulse width, while estimating the beta power (13-30 Hz) in the globus pallidus internus (GPi) for each DBS parameter. This process generated 200 distinct samples (**Fig. 1c** and **Supplementary Fig. 1**).

We obtained the second dataset from one non-human primate (NHP), a male Macaca fascicularis (CRP, Port Louis, Mauritius) weighing 12 kg. We referred this data as NHP data. During a preliminary period, the animal was progressively trained to remain on a primate chair in preparation for experimental sessions. The surgical procedure was carried out under general anesthesia after a fasting period of 10 hours. Initially, an intramuscular injection of 0.4 mg ketamine (Imalgene, Merial laboratory, France) was administered for induction, followed by a maintenance dose of 0.2 mg per hour. Additionally, 0.2 ml of 2% xylazine (Rompun, Bayer Healthcare AG, Germany) was given at induction, with a subsequent maintenance dose of 0.1 ml per hour. To further alleviate discomfort, local scalp anesthesia was provided through a subcutaneous injection of lidocaine chlorhydrate. Throughout the preoperative period, prophylactic measures were taken, including the administration of antibiotics, analgesics, and anti-inflammatory drugs to ensure the well-being of the subject.

The surgery was conducted using a stereotactic frame (David Kopf Instruments, Tujunga, USA) and was monitored through intraoperative tele-radiographic control. To localize the hippocampus, bi-commissural landmarks were obtained via ventriculography, which involved the injection of a 2 ml water-soluble iodine contrast medium (Bracco Imaging, France). For hippocampal stimulation, two quadri-electrode leads were implanted within the hippocampus. The lower contact of the Summit RC+S lead (E0) was positioned at a depth of 30 mm from the dura, with the lead length being 40 mm, electrode length 1.27 mm, and outer diameter of the electrode 1.5 mm, spaced 0.5 mm apart (DIXI, France). All components were secured to the skull with fixing screws and encapsulated in a dental acrylic cap. Preparation for the placement of the neurostimulator (Summit RC+S, Medtronic, Minneapolis, USA) involved creating a subcutaneous space in the back of the animal after making a skin incision. This device was capable of both stimulating targets and recording local field potentials (LFPs). Lead extensions (model 37087, 40 cm, Medtronic, Minneapolis, USA) were routed through the subcutaneous space in the back and neck and connected to the implanted electrode and stimulator. Post-surgery, the monkey was allowed to recover and was closely monitored. Food and water were provided ad libitum. All procedures were conducted in accordance with the European Communities Council Directive of 2010 (2010/63/UE) and the guidelines of the French National Committee (2013/113). The authorization (#00132.01) for conducting the experiments was granted by the Committee on the Ethics of Animal Experiments (#04). All experimenters were properly trained and certified for animal experimentation, ensuring that every effort was made to minimize animal suffering while maximizing the data obtained.

In our study, we utilized the Summit implantable neurostimulator (INS)^30,31^ for recording from and stimulating the hippocampus (**Fig. 2a**). To control the recording configuration, data streaming, and stimulation, we connected the INS to the research host computer (RHC) using the application programming interface (API) provided by the University of California, San Francisco (UCSF) (https://github.com/openmind-consortium/App-aDBS-ResearchFacingApp). We implemented an asynchronous distributed multielectrode stimulation (ADMES) approach in the Summit RC+S system to stimulate the NHP hippocampus. ADMES is a novel brain stimulation technique that has demonstrated effectiveness in reducing pathological synchronous activities and seizure frequency in a mesial temporal lobe epilepsy (MTLE) rodent model^14^. **Fig. 2b** illustrates a conceptual diagram of ADMES and its implementation in the Summit RC+S system. For the ADMES configuration, we utilized programming Group A of the Summit RC+S and leveraged the intrinsic delay across four programs in this group to create an asynchronous stimulation pattern. The post-stimulation signals were band-pass filtered between 0.5 and 80 Hz and sampled at a rate of 500 Hz. In this parameter sweep experiment, we varied the stimulation amplitude, frequency, and pulse width. We tested five amplitudes (100 µA, 200 µA, 300 µA, 400 µA, and 500 µA), ten frequencies (7 Hz, 12 Hz, 31 Hz, 50 Hz, 65 Hz, 72 Hz, 100 Hz, 130 Hz, 145 Hz, and 180 Hz), and two pulse widths (200 µs and 400 µs), resulting in 100 different combinations of the three stimulation parameters. Safe amplitude and frequency levels were established through an active discharge experiment. Each stimulation lasted 10 seconds, followed by a 15-second recording of the post-stimulation signal. The entire experiment took approximately 2500 seconds, or 42 minutes. We repeated this experiment three times on the same subject on the same day. During the experimental sessions, the NHP was positioned on a primate chair (Crist Instruments, USA) allowing free movement of limbs and oral feeding during the experiments. Penicillin salt (penicillin G sodium 3032, Sigma-Aldrich) was diluted with sterile water for injection (1000 IU / µL). Using sterile technique, a range from 3000 to 15000 IU of penicillin was injected at a rate of 2 to 3 µL / min with a Hamilton syringe and pump through a cannula, within the right HPC, at a depth of 28 mm from the dura. Each experiment corresponded to a single penicillin injection which was performed in the morning allowing monitoring throughout the day. A clinical epileptologist observed no seizure activity or after discharges during the parameter sweep experiment or in a thorough post-experiment review of all recorded LFP channels. Our previous study provided the details of this dataset ^32^.

For the third dataset (optogenetic data), we used one adult male Sprague-Dawley rat (2– 3-month-old; 250–300 g) from Charles River Laboratories (Wilmington, MA, USA). The animal was maintained within a 12/12 light/dark cycle vivarium while accessing unlimited food and water. Emory University’s Institute for Animal Care and Use Committee approved all procedures used in this study.

Two-step surgical procedures under anesthesia (1.5%–4% inhaled isoflurane) were performed as described in ^33^. In the first step, we injected the viral vector (AAV5-hSynapsin-Channelrhodopsin2-eYFP) into the medial septum just to the right of the midline at a 20° angle to the dorsal-ventral axis (0.40 mm anterior, 2.12 mm lateral at the 20° angles, 5.80 mm ventral to pia along the rotating axis). Using a pulled-glass pipette attached to a stereotactically mounted injector (Nanoject, Drummond Scientific Co., Broomall, PA, USA), we injected a volume of 1.8 µl containing 10^12^ particles ml^−1^ with a rate of 0.35 µl min^−1^. After the injection and drawing of the pipette, the scalp was stapled closed, and Meloxicam was administered as an analgesic (3–5 mg kg^−1^).

The second surgery step was conducted after two weeks, allowing time for recovery and optogenetic channel expression after the first one. We implanted a 16-channel multielectrode array (MEA; Tucker Davis Technologies (TDT), Alachua, FL., USA) in hippocampal CA3 and CA1(centered at 3.50 mm posterior and 2.80 mm lateral to bregma). The ferrule was then inserted at a 20-degree angle to the dorsal-ventral axis into the reopened initial injection craniectomy, approximately 5.8 mm from the pia along the rotating axis. For 10 seconds, a 17 Hz, 10 ms, 50 mW mm^2^ stimulation was used to identify the correct ferrule depth. Finally, dental acrylic was used to seal the craniectomy and keep the electrode and ferrule in place. Also, the electrode ground, reference wires, and structural support were fixed to the skull with five 2 mm stainless steel screws.

After the subject recovered from the second surgery, we conducted the parameter sweep experiment on the anesthetized animal. We started by placing the animal under 3.0% isoflurane gas by volume (carried by O_2_) and then waited for the animal’s breathing to stabilize at 1 breath per second. We adjusted the anesthesia when their breathing started to go too high or too low beyond 1/sec. The typical range for anesthesia was 1.5-3%. After stabilizing the depth of anesthesia, we delivered the light to the medial septum through the implanted fiber optic at all combinations of amplitude (between 10 to 50 mW mm^−2^), pulse-width (between 2 to 10 ms), and frequency (between 5 to 42 Hz) for a total of 72 combinations of stimulation parameters. Stimulation parameters were applied in a random order for 5 s. The LFP signals were recorded from the hippocampus throughout the experiment using an RZ2 BioAmp Processor and a PZ2 pre-amplifier TDT (Alachua, FL, USA). Signals were recorded at a sampling rate of 24414 Hz and then downsampled to 2000 Hz for further processing and to limit the computational load during real-time experiments.

### Preprocessing and spectral power estimation

When preprocessing the NHP and optogenetics data, we removed the very noisy signal, by a visual inspection, before filtering the signals with a low pass filter between 0-80Hz and a notch filter at 50 Hz for NHP and 60 Hz for the optogenetics dataset. In the NHP and optogenetics datasets, we calculated spectral power using the Thomson multi-taper method from the Chronux toolbox (http://chronux.org/), implemented in MATLAB (Mathworks, Natick, MA). The parameters for spectral analysis were the following: moving window = 5 s with 0.1 s overlap, time-bandwidth product (TW) = 3, number of tapers (K) = 5. For the NHP dataset, we estimated the post-stimulation theta power (5-8 Hz). While we could have used power from other frequency bands to test our framework, we focused on the theta band (5-8Hz) due to its correlation with seizures in MTLE, as reported in previous studies ^34,35^. For the optogenetics data, we estimated the during-stimulation (5 seconds) gamma power (31-55 Hz).

### In-silico modeling

**Fig. 1a** provides an overview of the *in silico* simulation process employed in our study to validate the AL framework. The process began with a parameter sweep experiment, where we varied the stimulation parameters within a safe range and estimated the brain response from the post-stimulation signal for electrical stimulation or during the stimulation signal for optogenetic stimulation. After obtaining the input matrix containing the stimulation parameters and their corresponding brain responses, we divided it into two sets: an unseen test dataset and a training dataset, which we refer to as the training pool dataset. The *in silico* modeling consisted of three phases: the *Initial Sampling Phase,* the *Model Improvement Phase,* and the *Validation Phase*. In the *Initial Sampling Phase*, we randomly selected a few samples (equal to the number of stimulation parameters) from the training pool dataset to form an initial training set and trained an initial model. In the *Model Improvement Phase*, we iteratively enhanced the model by adding one sample at a time to the training data from the training pool dataset. We employed both AL and RS approaches to refine the model in this phase and repeated this process *N* times. Within each iteration of the *Model Improvement Phase*, we validated the model on the unseen dataset, effectively running the *Validation Phase* within each iteration of the *Model Improvement Phase*. In each iteration of the *Model Improvement Phase*, we calculated the root mean square error (RMSE) between the actual and predicted brain responses on the unseen test dataset.

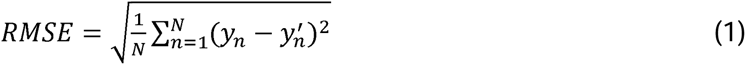

where *y_n_* and 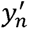 were the true and the predicted label (i.e., brain response). We continued the procedure until *i* = *N*. The output of this process was a curve similar to what is shown in **Fig. 1b**. The first and second curves represented the RMSE on the unseen test data in each iteration, respectively. At the end of the process, we calculated the area under the curve (AUC) for each RMSE curve. To obtain statistics, we ran the entire process 1000 times and compared the result of RS and AL with different query strategies. The query strategies included QBC, EMCM, greedy sampling (GS), RD, QBC+RD, EMCM+RD, GS+RD, enhanced QBC (EQBC), and enhanced EMCM (EEMCM). Additionally, we used ridge regression (RR), SVR (with both linear and Gaussian kernel functions), and Gaussian process regression (GPR) models to evaluate the utility of RS and AL with different query strategies.After calculating the AUC_RMSE for each model, we compared the AUC_RMSE of the nine AL models with that of RS using a two-sample t-test with the Bonferroni correction, setting the significance level at 0.05/9= 0.0056 due to the nine comparisons.

### Regression model

Depending on the linearity or non-linearity of the effect of stimulation, we could use either linear or nonlinear regression models. When assuming a linear relationship between the stimulation parameters and brain response and using linear regression, we needed to minimize the loss function in

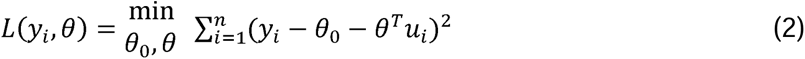

Where *θ* denoted the regression coefficients, *u_i_* contained the stimulation parameters, and *y_i_* was the brain response. In this study, we used ridge regression and support vector regression (SVR) with a linear kernel function as linear regression models to find the functional link between the stimulation parameters and the brain response. The ridge estimator solved the equation below:

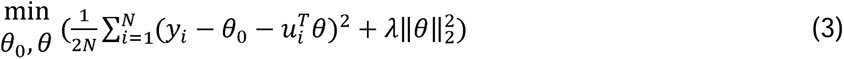

SVR tried to minimize

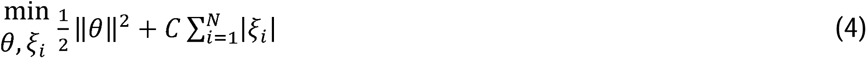

with the constraint

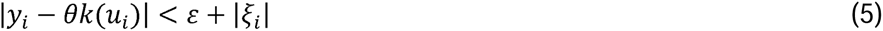

The SVR dual formulation was

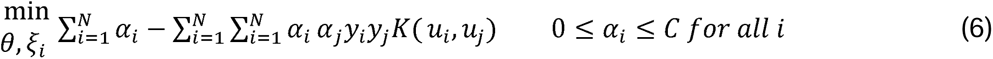

subject to

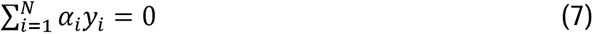

in which *k*(.) was the kernel function. In the linear SVR, we used the linear kernel function as shown below:

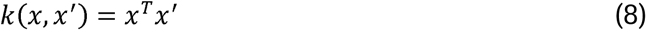

The nonlinear regression was an extended version of the linear one that used a much larger and general function class. This method estimated the function *f*(*u*; *θ*) as an additive expansion based on the basis function *b*(*u*; *β_m_*):

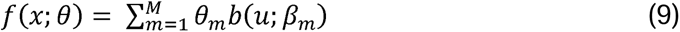

in which *β_m_* was the mean of split location and the terminal node for each splitting variable, *θ_m_* was the coefficient estimated by minimizing the loss function, which was *L*(*y*, *f*(*x*; *θ*)) = *y* − *f*(*x*; *θ*))^2^. In our study, as the nonlinear regression models between the DBS parameters and brain response, we used support vector regression with a radial basis function (RBF) kernel function in Equation (10).

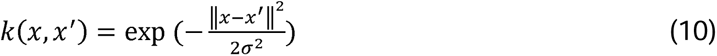

where exp(.) was the exponential function, ||.|| is the Euclidean norm for vectors, and *σ* was the kernel parameter determining the geometrical structure of the mapped samples in the kernel space. Additionally, we used Gaussian process regression (GPR) as another nonlinear regression model.

The GPR was written as:

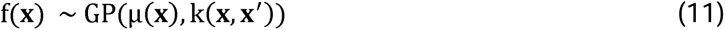

where μ(***x***) was the mean function, *k*(***x***, ***x′***) was the covariance function of the GP, and ***x***, ***x′*** were the input training data used to create the model. In GPR, we assumed that the training brain response followed a zero-mean prior Gaussian distribution.

### Query strategy

After building an initial model based on a few random stimulation parameters and their associated brain response, we used a query strategy to select the stimulation parameter most likely to improve the regression model. The query strategy was designed to meet three criteria: *informativeness*, *representativeness*, and *diversity*. *Informativeness* meant that the query strategy should select the stimulation parameters with the richest information (i.e., those parameters that improve the model more than the others). Previous studies showed that query by committee (QBC) and expected model change maximization (EMCM), which are based on uncertainty sampling, can query the samples with the richest information ^36^. The number of stimulation parameters in the vicinity of the selected stimulation parameter for the next query evaluated the *representativeness* of the selected parameter. By evaluating a selected parameter’s *representativeness*, we ensured that the selected parameters were not outliers. For example, the red circle “A” in **Supplementary Fig.6** does not meet the *representativeness* criterion since it is more likely to be an outlier. *Diversity* meant that the selected parameter sets should be distributed across the entire parameter space. By considering the selected parameter’s *diversity*, we ensured that the selected parameter was not from a small local region of the parameter space. For example, the red circle “B” in **Supplementary Fig.6** is the next best sample that would need to be collected if we were to meet the *diversity* criterion. A previous study introduced the representativeness and diversity (RD) method to meet both *representativeness* and *diversity* requirements in the query strategy. Our study uses query methods that meet all three criteria mentioned above ^37^.

### Query by committee

Query by committee (QBC) is a popular active learning query strategy in both classification ^38^ and regression ^39,40^. In contrast to uncertainty sampling, which uses a single model, QBC builds a committee of models (learners) from existing labeled training data, which in our case consists of stimulation parameter sets and their respective brain responses. QBC then selects the unlabeled data (i.e., unadministered stimulation parameters) that the committee most disagrees, i.e., sample by showing most deviation from mean, should be sampled in the next step. In this approach, we first bootstrapped the *S_0_*labeled sample into *p* copies, called *C_1_* to *C_p_* in **Supplementary Fig.7**, where each copy contained *S_0_*samples with duplication, and then for each copy, we made a regression model, called *M_1_* to *M_p_* in **Supplementary Fig.7** ^41^. Therefore, in total, we had *P* regression models. Next, for the *N-S_0_* unlabeled sample, we computed the variance of *P* predictions using the equation below:

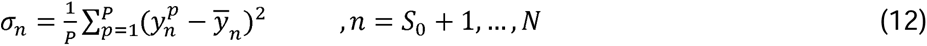

where 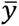 was the average of predicted values of an unlabeled sample from *p* models (i.e., 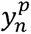) as shown in the equation below:

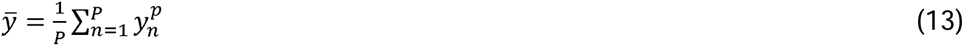

The sample with the highest *σ_n_* was selected for the next query.

### Expected model change maximization

Expected model change maximization (EMCM) is another popular method for querying samples for both regression ^42^ and classification ^43^. First, from the *M_0_* labeled samples, we built a regression model and computed the output of the model for unlabeled data (i.e., *x_n_* which we called 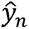. Similar to the QBC approach, we used the bootstrap method to construct *P* regression models from the labeled samples, and then we predicted the n^th^ unlabeled sample (i.e., 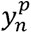) from each *P* model. Then we calculated the confidence of the prediction using the equation below:

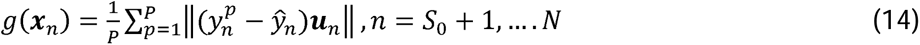

When querying the label in the next step, we selected the sample with the maximum *g*(***x****_n_*). **Supplementary Fig.8** illustrates the EMCM strategy.

### Greedy Sampling

In the GS method, we selected unlabeled samples (i.e., unadministered stimulation parameters) based on their distance from previously labeled samples (i.e., administered stimulation parameters) ^44^. For each unlabeled sample (i.e., *x_n_*) in the *N-S_0_* set, we calculated the distance between that sample and all of the samples in the previously labeled samples set (i.e., *u_m_*).

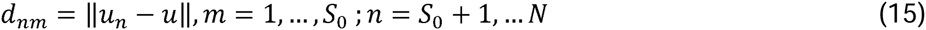

Next, we selected the unlabeled sample with the maximum distance from the labeled samples.

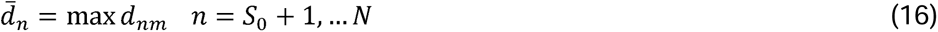

### Integrating representativeness and diversity (RD) with QBC, EMCM, and GS

The main drawback of the methods discussed above was that they did not consider the sampling *representativeness* and *diversity*. The RD approach added both *representativeness* and *diversity* to the approaches we discussed in the previous section. By adding these two criteria to QBC, EMCM, and GS, we hoped to improve the outcome by reducing the estimated error and increasing the correlation between the actual and predicted brain response on the unseen stimulation parameters, i.e., those parameters that had not been previously collected. To illustrate how this method worked, let’s assume that we had *d* labeled samples in our entire dataset (with the size of *N*). Therefore, we had *N - d* unlabeled samples. In the naïve RD algorithm, we first performed k-means clustering on the entire dataset, where *k=d+1*. Since we had only *d* labeled samples, at least one cluster did not contain any labeled samples. However, in practice, some clusters might have had more than one labeled sample; therefore, we should have had more than one cluster without any labeled samples.

Next, to meet the *diversity* requirement, we needed to select the largest cluster that contained no labeled samples for the next query. To meet the *representativeness* requirement, the selected sample needed to be the sample closest to the center of the cluster. In the integrated version of RD with QBC, EMCM, and GS, after selecting the largest cluster without a labeled sample, we selected the next sample for labeling based on QBC, EMCM, and GS on all samples in the selected cluster. An illustration of this method is shown in **Supplementary Fig.9** where we wanted to model a link between the stimulation parameter (x axis) and the target brain response biomarker (y axis). We had two know samples (the black circles), and the unknown samples were shown by gray circles. Since *d*=2 (the known samples), we separated the entire parameter space into three clusters, where we would have at least one cluster without any known (black) samples. This cluster was shown in a red ellipse in **Supplementary Fig.9**.

### Enhanced QBC and EMCM

The initial, randomly selected samples can sometimes be outliers, and their selection can reduce regression model performance. Conventional QBC and EMCM approaches do not involve any mechanisms that prevent the selection of outliers. As such, to prevent the effects of outlier selection, we used the EQBC and EEMCM methods that were proposed in ^18^.

### Brain state dynamic modeling

The modeling of the effects of different stimulation parameters on target neurophysiological brain response biomarkers can be formulated as a dynamic or static mapping. In the dynamic formulation, predicting the effect of stimulation is a function of time and the current neural state (**Fig. 3a**). In traditional static models, time is ignored and thus possibly confounds the effects of the variables being studied. In dynamic models, time is fundamental to both the basic structure of the data and the understanding of how a process unfolds. The mathematical description of the dynamic model of the brain can be expressed by a set of *n* coupled first-order ordinary differential equations called state equations.

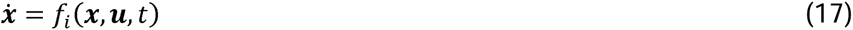

In this set of equations, ***x***=[*x_1_,x_2_,…,x_n_*] is the state vector, ***u***=[*u_1_, u_2_, …,u_r_*] is the input vector, here are the stimulation parameters and *f(.)* is a vector function, here is the regression model. Equation below shows a linear representation of this model.

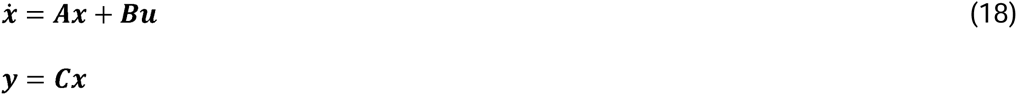

In this equation, ***x*** ∈ ℝ*^n^* is the state vector, ***ẋ*** ∈ ℝ*^n^* is the state derivative, and ***y*** ∈ ℝ*^p^* is the output vector. ***A*** ∈ ℝ*^n×n^* is the state transition matrix, ***B*** ∈ ℝ*^n×m^* is the input and ***C*** ∈ ℝ*^p×n^* is the output matrix ^45^. As mentioned earlier, the incorporation of the brain state prior to stimulation in the dynamic model was the only difference between the dynamic and static models. With this new perspective, we updated our *in silico* modeling process by adding the state of the brain before stimulation.

### Stochastic response dynamic modeling

Unlike other systems, the brain’s response to a specific input is somewhat stochastic, meaning that its response is not always consistent across multiple system identification tests. In practical terms, this means that identical stimulation parameters applied at different times can result in varying brain responses (**Fig. 3b**). To capture the dynamics of the brain, we updated our original framework to incorporate multiple responses of the same parameter set collected across multiple sessions of the parameter sweep experiment (**Supplementary Fig.10**). This revised process, similar to the original framework shown in **Fig. 1a**, consists of three phases. Initially, we divided the entire dataset (i.e., sets of stimulation parameters and their associated brain responses) into an unseen validation set and a training pool dataset. In Phase 1, we randomly selected three stimulation parameters and one associated brain response for each parameter (from the *m* available responses). In Phase 2, we iteratively expanded the training dataset through *N* iterations. Specifically, based on recommendations from either the AL or RS approach, we selected one stimulation parameter and its associated brain response (from all *m* available samples) to update the model. After each model update, we validated the model on the 20% unseen validation dataset and calculated the RMSE between the predicted and actual brain responses (Phase 3). We then computed the AUC for the RMSE curve at the end of the process. We repeated the entire process 1000 times to gather statistical data. Finally, we compared the mean AUC_RMSE of the RS model with those of the AL models using all query strategies.

### *In vivo* experiment

After finding the best query strategy via *in silico* modeling, we compared AL and RS through an *in vivo* test (**Fig. 5a**). Before starting the experimental session, we put the entire parameter set, including stimulation intensity, frequency, and pulse width, into the unseen test dataset (25 samples) and the training pool dataset (1025 samples). Then, we selected three sets of stimulation parameters in the first phase, applied optogenetic stimulation to the medial septum of the anesthetized rat, recorded the CA1 signal during stimulation, and estimated the slow-gamma power (30 to 55Hz). Using the best regression model obtained through the *in silico* modeling, we trained a regression model between the stimulation parameters and slow-gamma power using these three sets of parameters. In Phase 2, we iteratively added *N* = 72 training samples one at a time in an analysis using RS and again in an analysis using AL. In Phase 3, we collected the brain response and estimated the slow-gamma power for the unseen test dataset. Each session took two hours. We repeated the entire two-hour experiment twenty times to generate statistics, in which we randomly alternated the order of RS and AL in Phase 2. Also, we used the same unseen test stimulation parameters in all twenty experiments. In the post-experiment analysis, we estimated the RS and AL model performance by calculating the RMSE on the unseen test data in each iteration of Phase 2, and we next estimated the mean AUC_RMSE of each experiment for both AL and RS models. We used the paired t-test because the same unseen test stimulation parameters were applied in each experiment, making the observations from the AL and RS models naturally paired and allowing for a more accurate comparison of their performance.

## Data availability

The data that support the findings of this work are available in the Supplementary Data files and have been deposited at: https://github.com/msendi6/active-learning-stimulation

## Code availability

The codes for training the machine learning models, analyzing the models, and generating the figures are available in GitHub link https://github.com/msendi6/active-learning-stimulation

## Author contributions

Conceptualization: M.S.E.S Methodology: M.S.E.S and E.R.C Investigation: M.S.E.S Analysis: M.S.E.S Visualization: M.S.E.S Writing –original draft: M.S.E.S and C.A.E Writing – review and editing: M.S.E.S, E.R.C, B.P, C.A.E, T.E.E, N.G.L, B.M, C.A.G, A.D, H.M, R.E.G, and V.D.C Supervision: H.M, R.E.G, and V.D.C, Funding acquisition: B.M, C.G, A.D, R.E.G, and V.D.C.

## Competing interests

Mohammad Sendi provides consulting services for Niji Corp. Helen Mayberg receives IP licensing fees from Abbott Labs and consults Abbott Labs, BlackRock Neuro, NextSense and Cogwear. Robert E. Gross serves as a consultant to Medtronic, which manufactures products related to the research described in this manuscript and receives compensation for these services. He also receives support for unrelated research.

## Funding/Support

This work was supported by The National Institutes of Health National Institute of Neurological Disorders and Stroke grant UG3-NS100559 (A.D, C.A.G, B.M, R.E.G) and R01MH123610 (V.D.C). Time spent preparing the revised manuscript for publication was supported by T32MH125786 (to W. Carlezon/K. Ressler, MPIs).

## Notes

https://github.com/msendi6/active-learning-stimulation

