## Supplementary information for "Refining Brain Stimulation Therapies: An Active Learning Approach to Personalization"

### **Cortex- Basal Ganglia-Thalamus computational model**

We used the biophysical model of the cortex- basal ganglia-thalamus network in the 6-OHDA lesioned rat of Parkinson’s disease proposed in Kumaravelu et al ^1^ to generate the synthetic data and validate the AL framework. This network contains the cortex, striatum, subthalamic Huxley type neurons. This model is shown in **Supplementary Fig.1**, and the model parameters are provided in Kumaravelu et al ^1^. In this model, *rCortex* neuron receives excitatory input from one Th neuron and inhibitory input from four randomly selected *rCortex* neurons. Moreover, each *iCortex* neuron receives excitatory input from four randomly selected *rCortex* neurons. Each *dStr* neuron receives excitatory input from one *rCortex* neuron and inhibitory axonal collaterals from three randomly selected *dStr* neurons. Each *dStr* neuron receives excitatory input from one *rCortex* neuron and inhibitory axonal collaterals from three randomly selected *dStr* neurons. Each *idStr* neuron receives excitatory input from one *rCortex* neuron and inhibitory axonal collaterals from four randomly selected *idStr* neurons. Each STN neuron receives inhibitory input from two *GPe* neurons and excitatory input from two *rCortex* neurons. Each *GPe* neuron receives inhibitory axonal collaterals from any two other *GPe* neurons and inhibitory input from all *idStr* neurons. Each *GPi* neuron receives inhibitory input from two *GPe* neurons and inhibitory input from all *dStr* neurons. Some *GPe/GPi* neurons receive excitatory input from two *STN* neurons, while others do not. Each *Th* neuron receives inhibitory input from one *GPi* neuron. To have healthy control (HC) and PD model, we change the M-type potassium current for both the direct and indirect medium spiny neurons (MSNs) by setting the maximal conductance at 2.6 mS/cm^2^ and 1.5 mS/cm^2^, respectively.

Using this model, we ran parameters sweep experiments for both HC and PD models, in which we estimated STN and measured the GPi beta power (13-30Hz). We developed a model to capture the functional relationship between the STN stimulation parameters and the changes in the GPi beta power (13-30 Hz). In more detail, we swept the stimulation frequency from 0 to 200 Hz with a 10 Hz increment and amplitude from 0 to 3 Am with a 0.2 mA increment and calculated the GPi beta power for each stimulation in both HC and PD patients. We generated data for both PD and HC models, as **Supplementary Fig.2** shows.

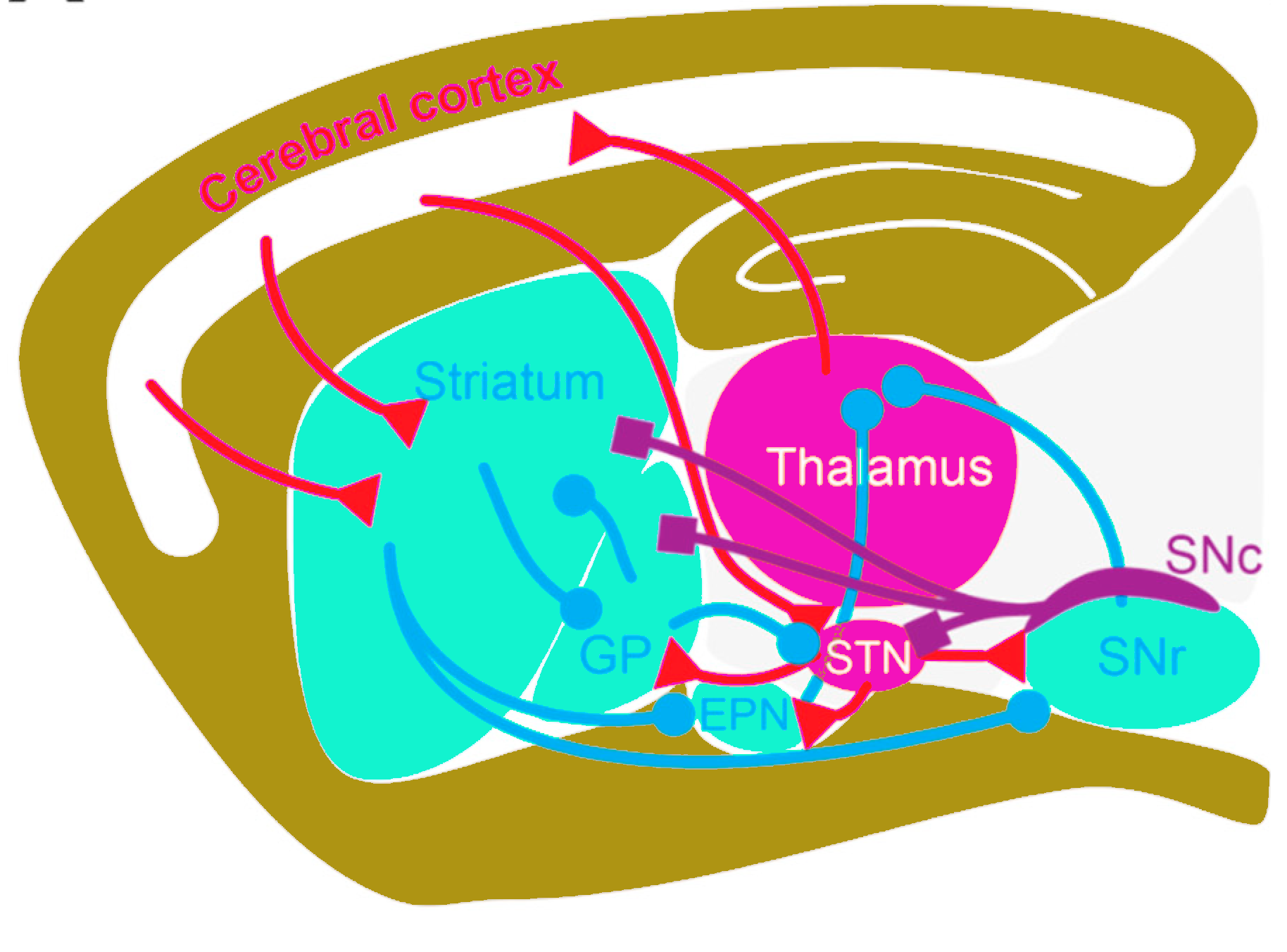

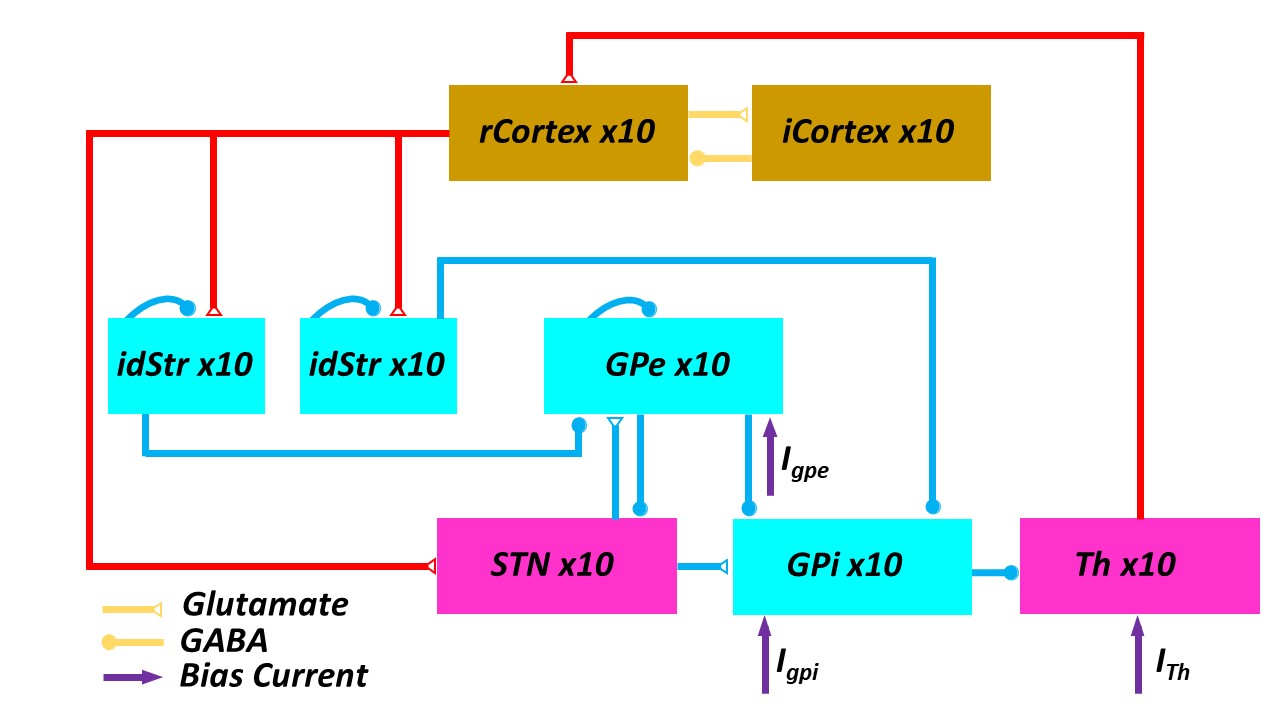

**a**

**b**

**Supplementary Fig.1. The PD model used to generate synthetic dataset. a)** The cortical-basal ganglia-thalamus in rat, **b)** The cortical-basal ganglia-thalamus network that shows the synaptic connection within the network model.

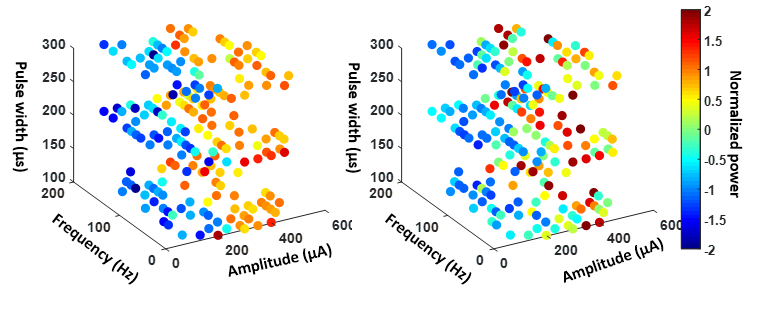

**a**

**b**

**Supplementary Fig.2. The synthetic data generated for cortical-basal ganglia-thalamus network. a)** PD model**, b)** HC model

**Supplementary Fig.3.** **Support vector regression (SVR) model with linear kernel function based on RS and AL with different query strategies on the cortex- basal ganglia-thalamus network HC model.** Mean RMSE_AUC of running 1000 times with different AL query strategies and RS sampling. Asterisks represent those AL strategies having significantly less AUC_RMSE than RS after Bonferroni corrections (corrected *p*<0.0056).

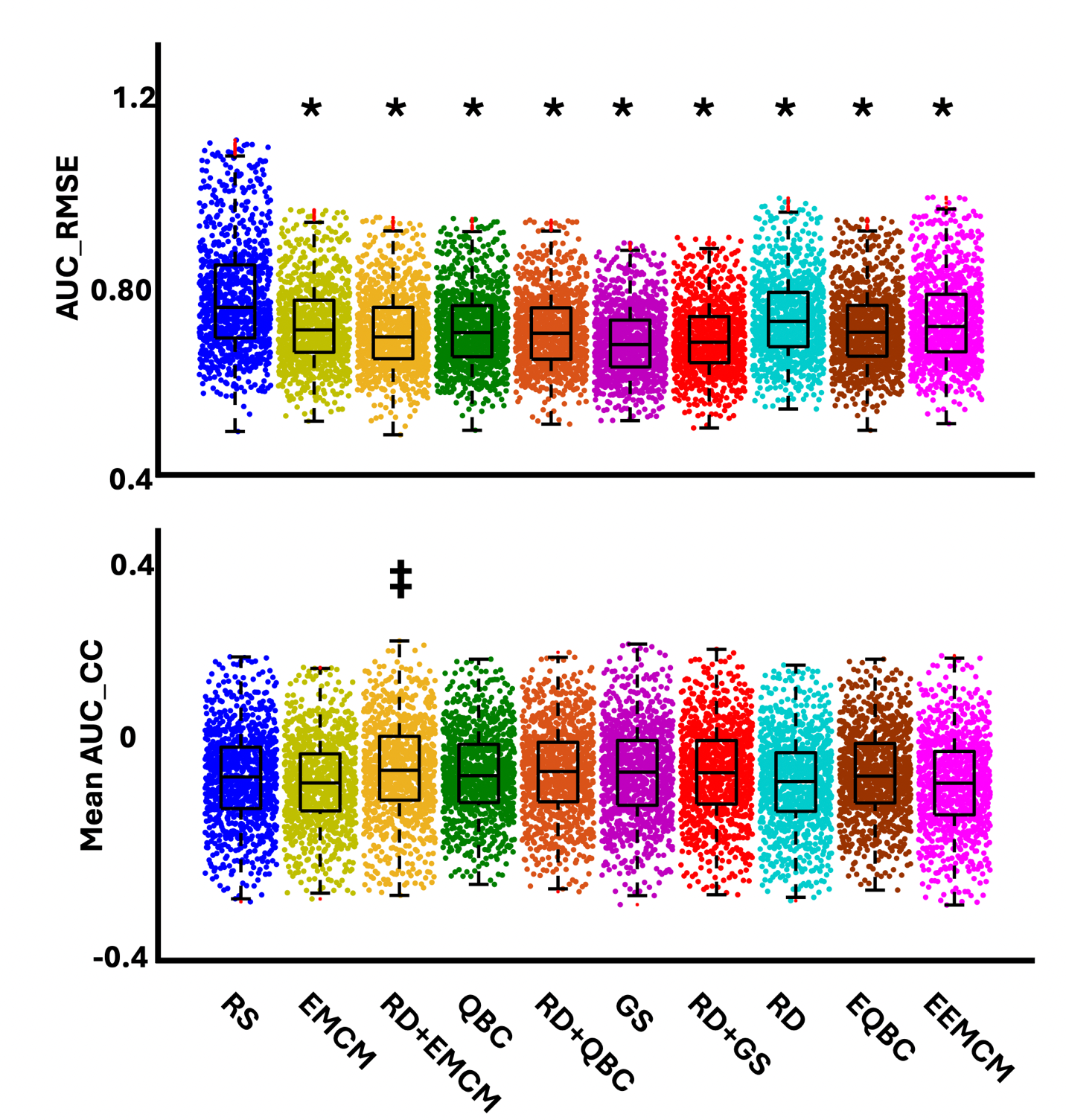

**Supplementary Fig.4.** **Support vector regression (SVR) model with Gaussian kernel function based on RS and AL with different query strategies on the cortex- basal ganglia-thalamus network HC model.** Mean RMSE_AUC of running 1000 times with different AL query strategies and RS sampling. Asterisks represent those AL strategies having significantly less AUC_RMSE than RS after Bonferroni corrections (corrected *p*<0.0056).

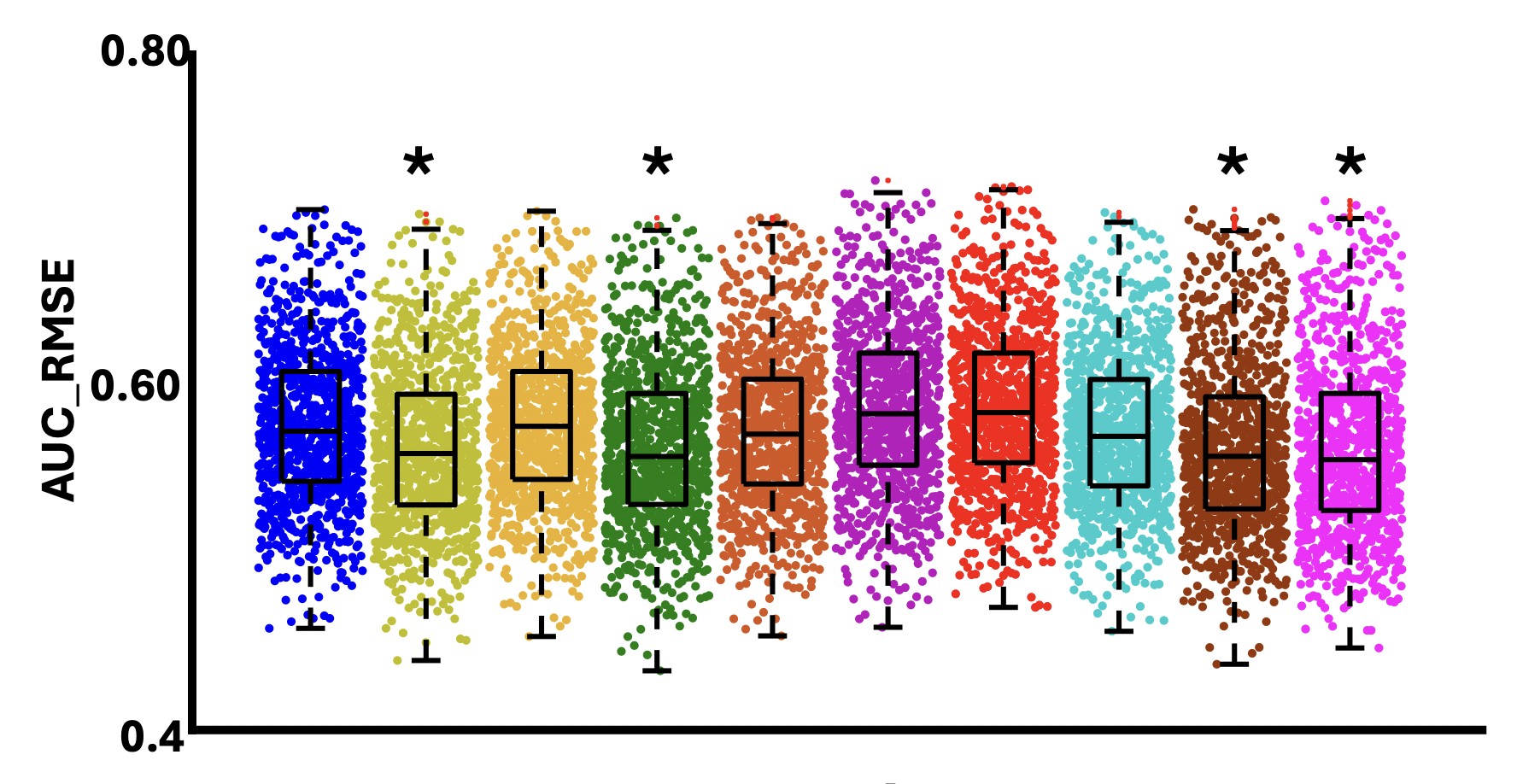

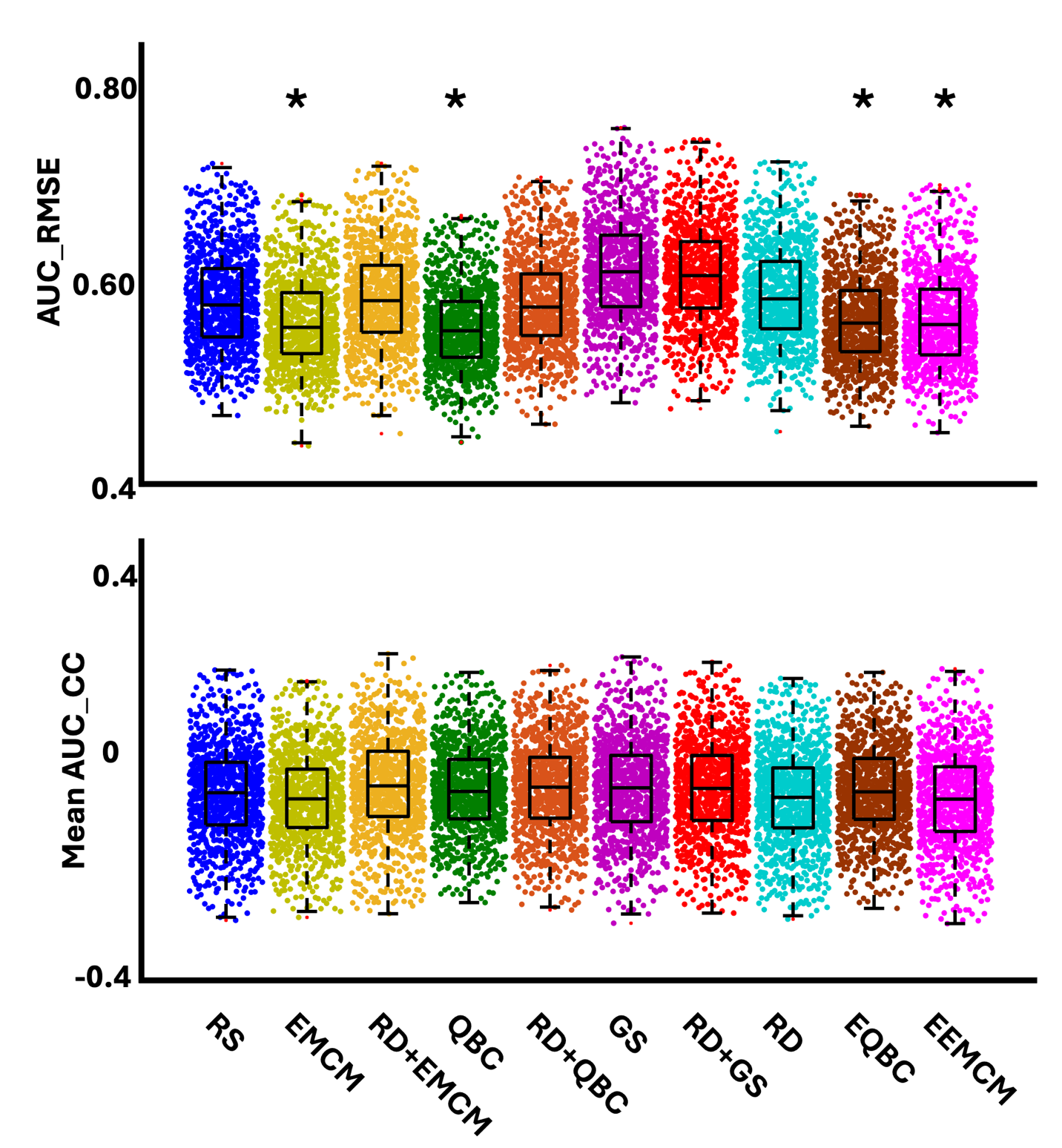

**Supplementary Fig.5.** **Gaussian Process regression model based on RS and AL with different query strategies on the cortex- basal ganglia-thalamus network HC model.** Mean RMSE_AUC of running 1000 times with different AL query strategies and RS sampling. Asterisks represent those AL strategies having significantly less AUC_RMSE than RS after Bonferroni corrections (corrected *p*<0.0056).

b

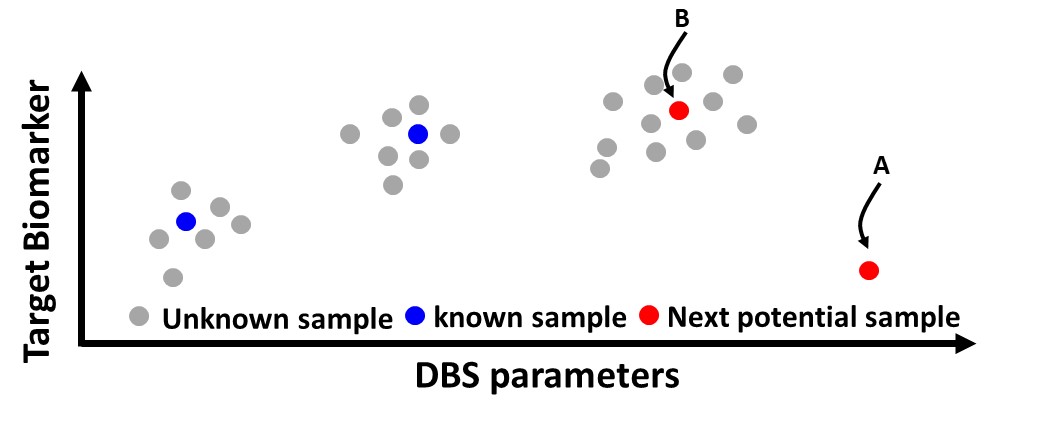

Supplementary Fig.6. Illustration of representative and diversity (RD) in active learning regression (ALR). The red circle “A” does not meet the representative criteria since it is an outlier. The red circle “B” does meet the diversity criteria.

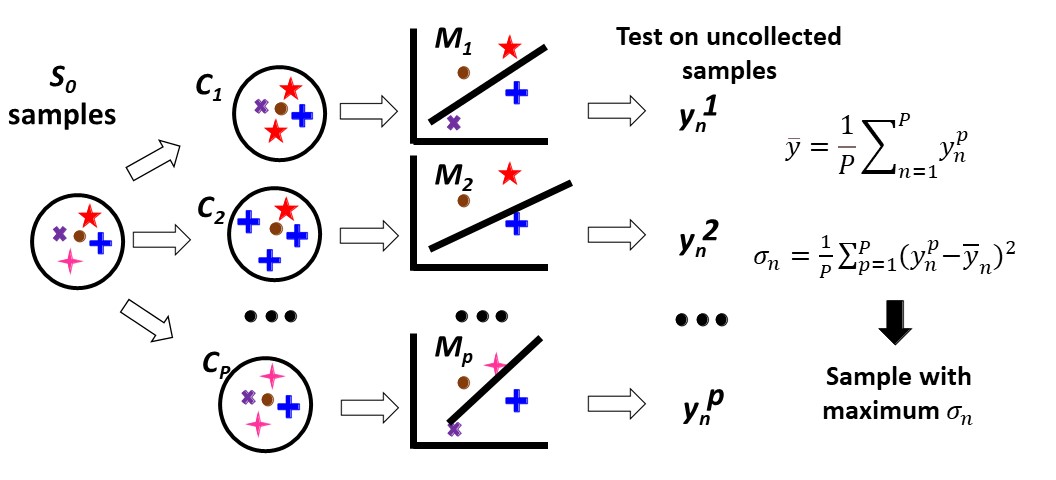

**Supplementary Fig.7.** Illustration of query by committee (QBC) in active learning regression (ALR).

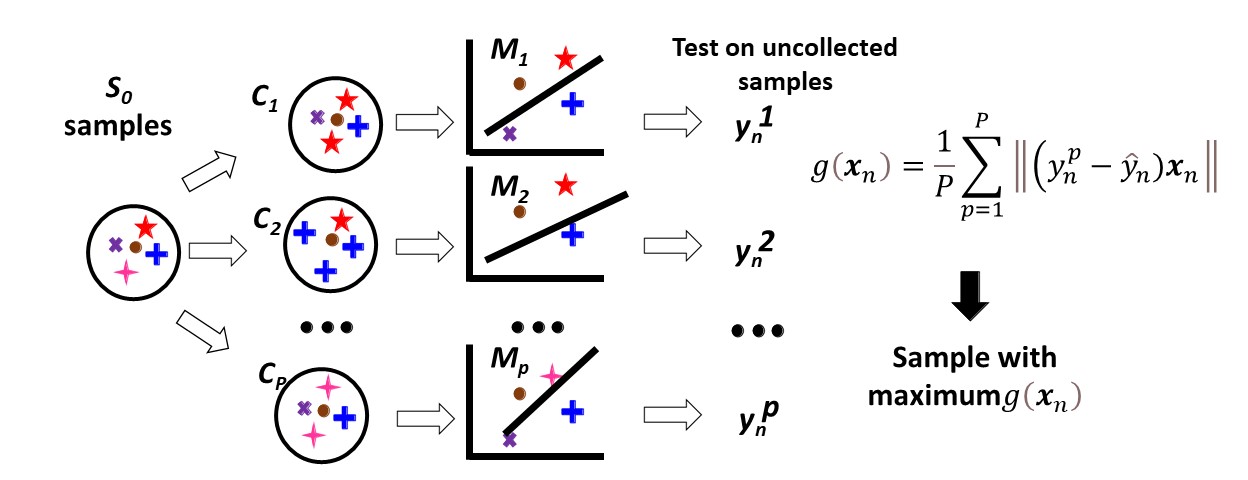

**Supplementary Fig.8.** Illustration of Expected model change maximization (EMCM) in active learning regression (ALR).

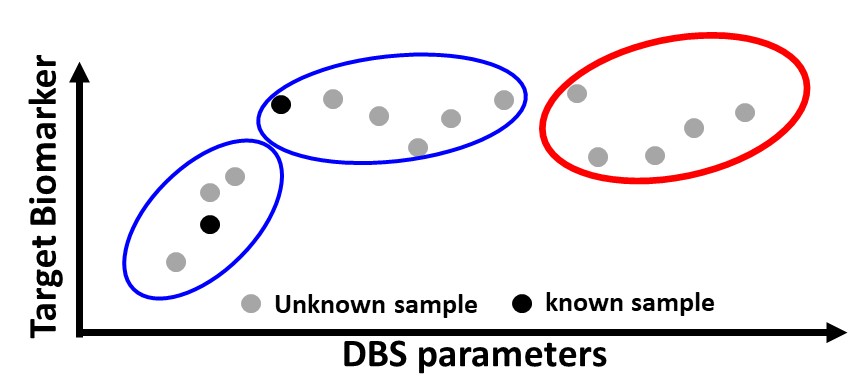

**Supplementary Fig.9.** Illustration of representative and diversity (RD) query strategy.

**Supplementary Fig.10. An overview of proposed stimulation for modeling the brain dynamics in the AL framework.** Same as the original framework, the simulation contains three phases. The only difference is the number of brain responses available for any specific stimulation parameter set. First, we put the entire available dataset into two unseen test datasets and a pool training dataset. In Phase1, we collect a few samples from the training pool dataset and build a model between the post-stimulation signal feature and stimulation parameters. We randomly select the brain response from the pool training dataset out of *m* available samples associated with selected stimulation parameters. In Phase 2, we iteratively update the model by querying a new sample. We would collect *N* samples, where the brain responses are selected randomly from *m* available samples associated with the selected parameters in the training pool dataset. At each iteration of Phase 2, we test the model with the unseen validation set and calculate the root mean squared error and the correlation between the actual response biomarker and the predicted value. Finally, we calculate the area under the curve or AUC of RMSE and correlation resulting from both AL- and RS-based models. We run the entire process 1000 times to get an statistics and finally we compare AL and RS model results with the ANOVA test.

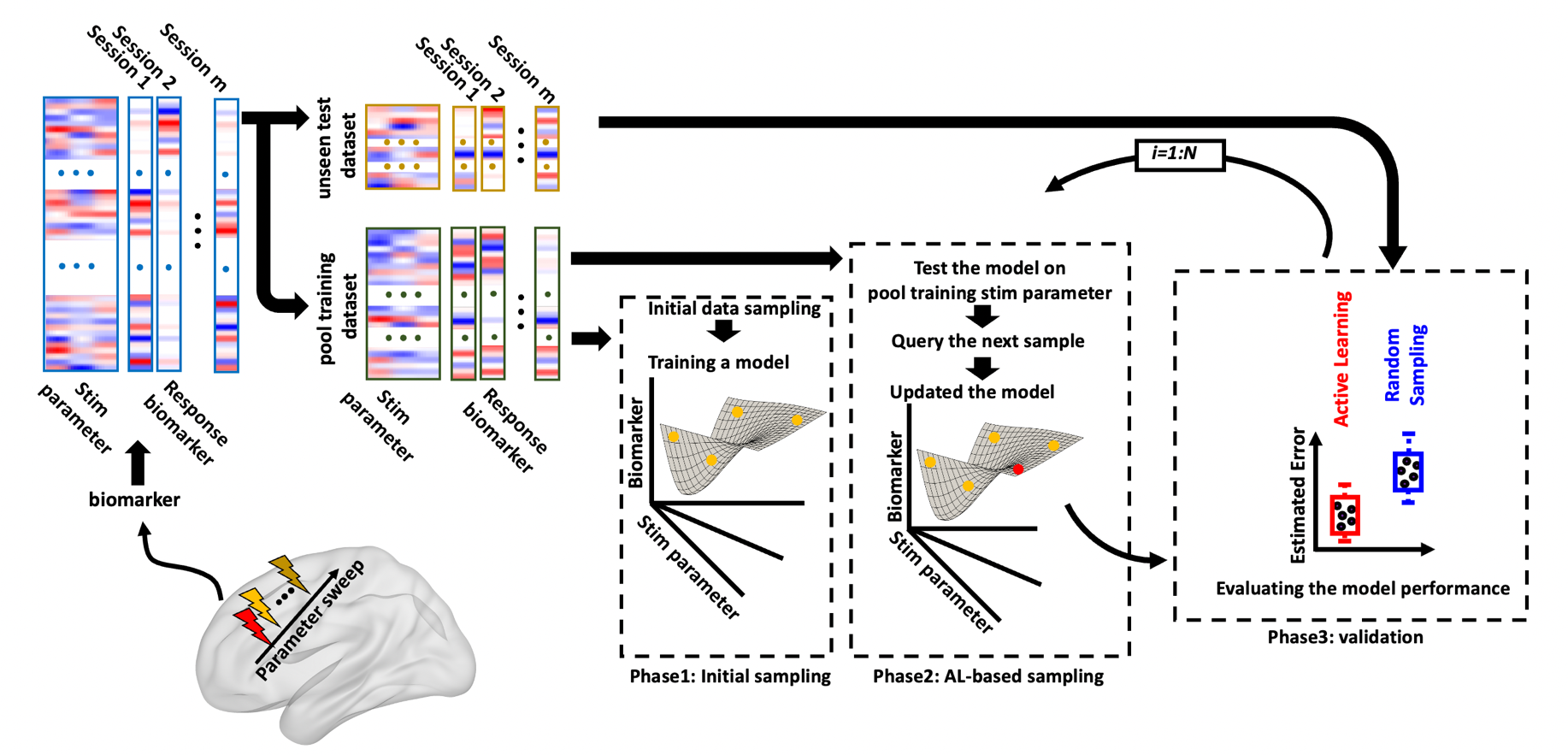
